# Viral proteins activate PARIS-mediated tRNA degradation and viral tRNAs rescue infection

**DOI:** 10.1101/2024.01.02.573894

**Authors:** Nathaniel Burman, Svetlana Belukhina, Florence Depardieu, Royce A. Wilkinson, Mikhail Skutel, Andrew Santiago-Frangos, Ava B. Graham, Alexei Livenskyi, Anna Chechenina, Natalia Morozova, Trevor Zahl, William S. Henriques, Murat Buyukyoruk, Christophe Rouillon, Lena Shyrokova, Tatsuaki Kurata, Vasili Hauryliuk, Konstantin Severinov, Justine Groseille, Agnès Thierry, Romain Koszul, Florian Tesson, Aude Bernheim, David Bikard, Blake Wiedenheft, Artem Isaev

## Abstract

Viruses compete with each other for limited cellular resources, and some viruses deliver defense mechanisms that protect the host from competing genetic parasites. PARIS is a defense system, often encoded in viral genomes, that is composed of a 53 kDa ABC ATPase (AriA) and a 35 kDa TOPRIM nuclease (AriB). Here we show that AriA and AriB assemble into a 425 kDa supramolecular immune complex. We use cryo-EM to determine the structure of this complex which explains how six molecules of AriA assemble into a propeller-shaped scaffold that coordinates three subunits of AriB. ATP-dependent detection of foreign proteins triggers the release of AriB, which assembles into a homodimeric nuclease that blocks infection by cleaving the host tRNA^Lys^. Phage T5 subverts PARIS immunity through expression of a tRNA^Lys^ variant that prevents PARIS-mediated cleavage, and thereby restores viral infection. Collectively, these data explain how AriA functions as an ATP-dependent sensor that detects viral proteins and activates the AriB toxin. PARIS is one of an emerging set of immune systems that form macromolecular complexes for the recognition of foreign proteins, rather than foreign nucleic acids.

## Introduction

Anti-viral defense systems in bacteria and archaea are extraordinarily diverse and many of these systems are mechanistically similar to immune responses in eukaryotic cells^1^. The recent expansion of bacterial and archaeal immune systems stems from the appreciation that defense systems tend to co-localize in the genome^2^ and that most systems are carried by mobile genetic elements, including prophages and satellite viruses^3^. The Phage Anti-Restriction Induced System (PARIS) is one of several defense systems recently identified within a hotspot of genetic diversity carried by P4 phage satellites^4^. The PARIS system of *Escherichia coli* strain B185 protects against phage T7 infection through a mechanism triggered by sensing the Ocr (overcoming classical restriction) protein of T7. Ocr is a DNA mimic that inhibits Type I Restriction-Modification and BREX defense systems^5,6^. Thus, PARIS is an ’anti-anti-restriction’ system that senses Ocr, a viral immune suppressor.

The PARIS defense system consists of two proteins, AriA and AriB (**Fig 1A,B**). AriA contains a conserved ABC ATPase domain, while AriB is a domain of unknown function (DUF4435), which includes a TOPRIM nuclease^7,8^. AriA and AriB are usually separate proteins, though they are occasionally fused into a single polypeptide. Several defense systems share an ABC ATPase domain and a TOPRIM domain, including the OLD (Overcoming Lysogenization Defect) protein produced by the P2 prophage, which blocks infection by phage lambda^9^. OLD does not directly sense a phage protein but rather the inactivation of the RecBCD exonuclease^10^. Activation of OLD results in translation inhibition^11^, but the underlying mechanism has not been determined.

**Figure 1:**
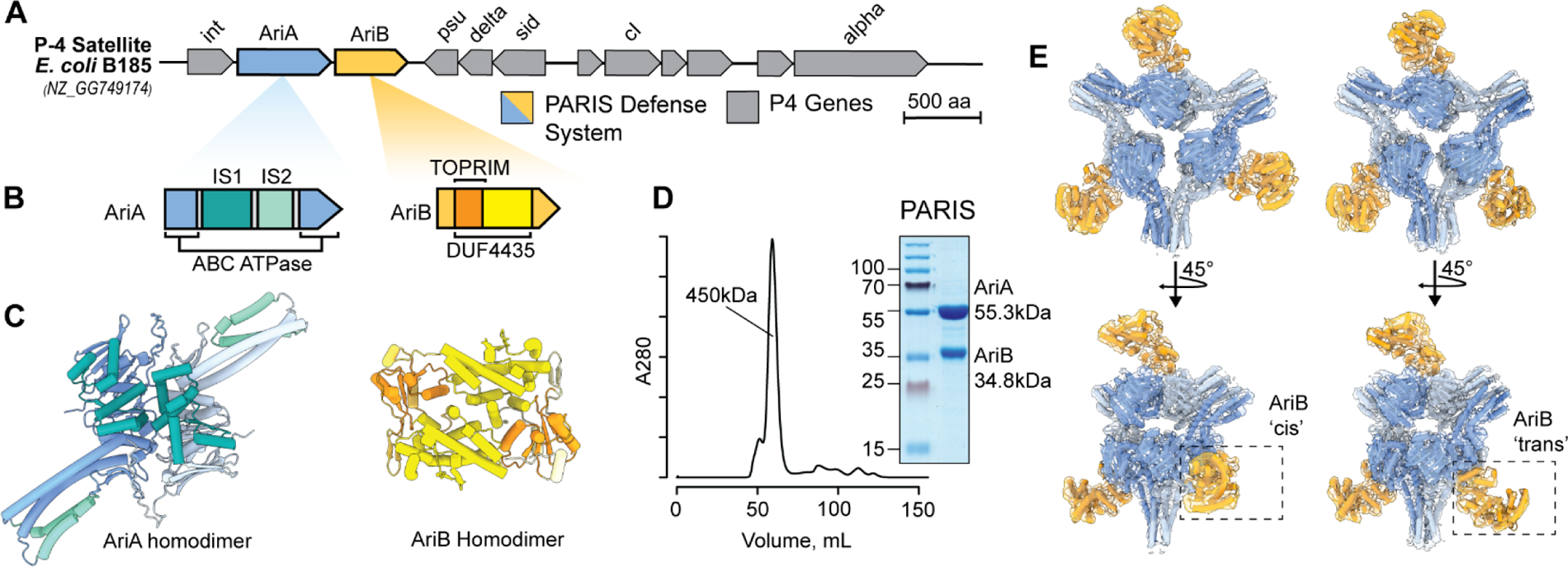
PARIS is a two-component system that assembles into a propeller-shaped supramolecular complex. A) Schematic of the integrated P4 satellite in the *E. coli* genome. B) PARIS genes colored according to domain organization. AriA is an ABC ATPase, while AriB a DUF4435 that includes TOPRIM nuclease. Insertion sequences 1 (IS1) is composed of three helices flanked by the ABC ATPase domain, while Insertion sequences 2 (IS2) adds two additional helices to the coiled-coil resulting in a helical bundle. C) AlphaFold2-predicted structures of AriA and AriB as homodimers colored according to structural features shown in panel B. D) Size exclusion chromatography reveals that AriA and AriB copurify as an ∼450 kDa complex. E) Experimentally determined density maps of the PARIS complex reveals a homohexameric assembly of AriA with D3 symmetry. The AriA hexamer is decorated by AriB subunits that attach to AriA homodimers in one of two possible orientations, a ‘*cis*’ arrangement where the three AriB subunits are facing the same direction, or a ‘*trans*’ arrangement where one AriB subunits is rotated by 180° relative to the other two.

Association between ABC ATPase and TOPRIM domains is an emerging theme in recently discovered immune systems^12^. Examples include the Gabija system^13,14^, effector toxin proteins of retron immunity^15^, and *bona fide* toxin-antitoxin (TA) modules^16^. This implies a broad adaptability of the conserved ABC ATPase and TOPRIM nuclease architecture to a variety of antigenic stimuli, resulting in the diversification of the immune modules likely acting through “abortive” mechanisms. We currently have little understanding of the molecular mechanisms underpinning how these systems sense and respond to viral infections.

Here we use cryo-EM to show that *E. coli* B185 PARIS systems assembles into a 425 kDa propeller-shaped complex with six molecules of AriA and three subunits of AriB. We show that AriA senses the T7 Ocr anti-restriction protein leading to the release of the AriB nuclease effector. Release of AriB from the AriA scaffold is necessary to potentiate AriB, which forms a nuclease-active dimer that degrades host lysine tRNA (tRNA^Lys^), thereby blocking translation and leading to growth arrest or cell death. We find that phage T5 carries its own tRNA^Lys^ isoacceptor, which rescue the phage from PARIS-mediated defense. Finally, we perform a phylogenetic analysis of PARIS systems, which provides insights into their evolutionary history and shows how they relate to other defense systems that employ ABC ATPases.

## Results

### PARIS assembles into a propeller shaped macromolecular complex

The PARIS defense system consists of two proteins, AriA and AriB, which protect cells from phages through a previously undetermined mechanism^4^. Here, we focused on the system from the P4 satellite integrated into the *E. coli* (str. B185) genome (**Fig 1A,B**). Based on published experimental structures of related proteins^8,13^ and AlphaFold2^17^ predictions (**Fig 1C, Sup Fig 1A-D**), we anticipated that AriA:AriB would assemble into a multi-subunit immune complex. To determine if and how AriA and AriB assemble, we overexpressed AriA with a C-terminally tagged AriB (AriB-Strep), pulled down on AriB, and purified the complex using size exclusion chromatography (**Fig 1D**). The AriA and AriB proteins co-elute in a single peak with an estimated molecular weight of 450 kDa. Next, we used cryo-EM to determine structures of the PARIS complex (**Fig 1E**). The structure explains how six molecules of AriA come together to form a D3 symmetric scaffold decorated by three subunits of AriB. The helical domains, which typically provide a platform for DNA binding in related ABC ATPases^18^, instead function as additional dimerization interfaces that enable a trimeric assembly of AriA homodimers (**Fig 1C, Sup Fig 2**). Dimerization of the helical domains from adjacent AriA homodimers form three blades of the propeller (**Fig 1E, Sup Fig 2)**. Additional density is evident between each of the AriA blades, where AriB attaches to AriA near the ATP binding sites. However, density in this region of the reconstruction reveals conformational heterogeneity, consistent with AriB attaching to either, but not both AriA molecules at the dimer interface (**Sup Fig 1E, Sup Movie 1**). Using multiple rounds of focused 3-D classification, we resolved two distinct isomers of the complex (**Fig 1E, Sup Fig 3D**). In one reconstruction, AriB attaches to three symmetrically equivalent AriA molecules (3.8 Å-resolution, *cis* configuration), while in the other reconstruction, one of the three AriB molecules binds the opposing AriA, which flips AriB 180-degrees relative to the other two AriB subunits (4.0 Å-resolution, *trans* configuration) (**Sup Fig 3, Sup Movie 1**).

**Figure 2:**
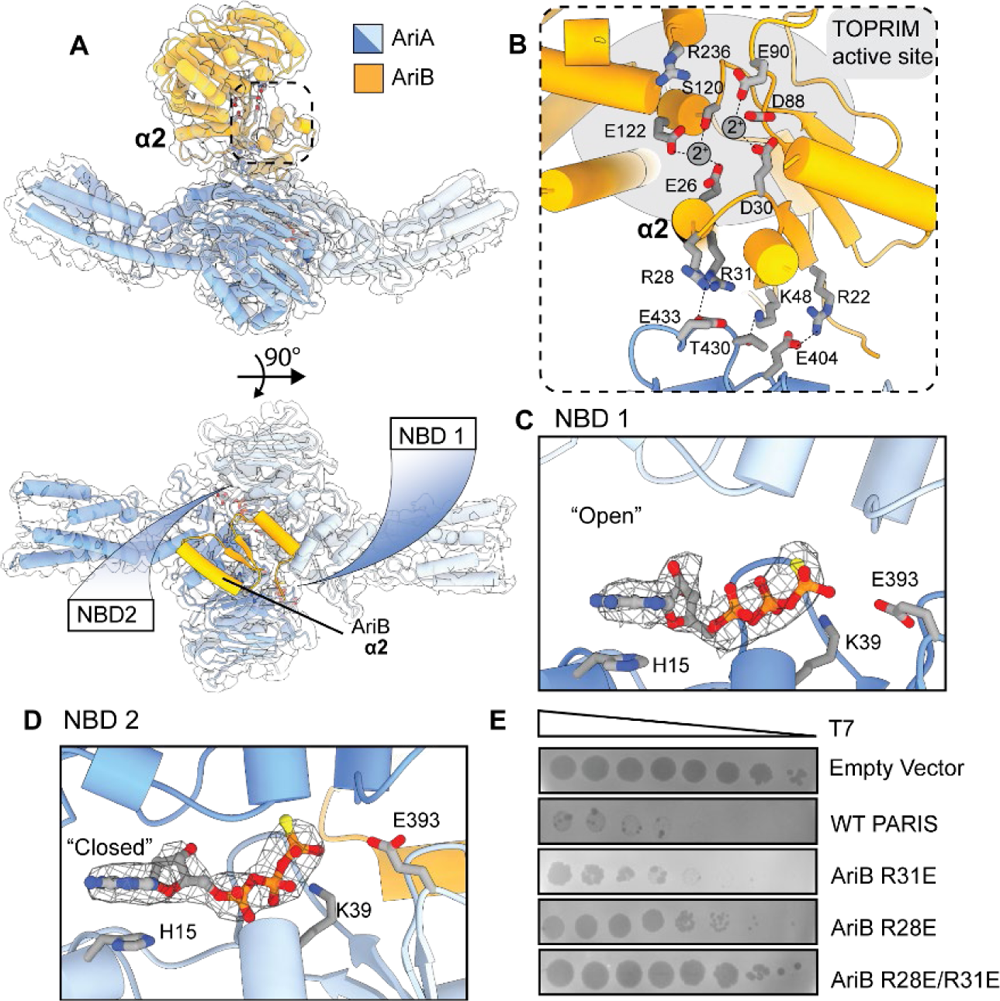
Interaction between the AriA ATPase and AriB are essential for PARIS-mediated defense. A) Each asymmetric unit of the PARIS complex contains an AriA homodimer with two nucleotide binding domains (NBD). A 90-degree rotation of the asymmetric unit, showing the N-terminus (res 1-53, yellow) of AriB positioned over the ATPase active sites of AriA. B) AriB attaches to AriA through electrostatic interactions near helix α2 of AriB. TOPRIM active site residues predicted to participate in metal binding (gray spheres) and catalysis are shown as sticks. C) Close-up of NBD1 showing ATPγS bound in an “open” conformation. D) Close-up of NBD2 showing bound ligand trapped in a “closed” pre-cleavage state. E) Plaque assays performed using a serial dilution of phage T7 on *E. coli* MG1655 carrying the WT or mutated PARIS system.

**Figure 3:**
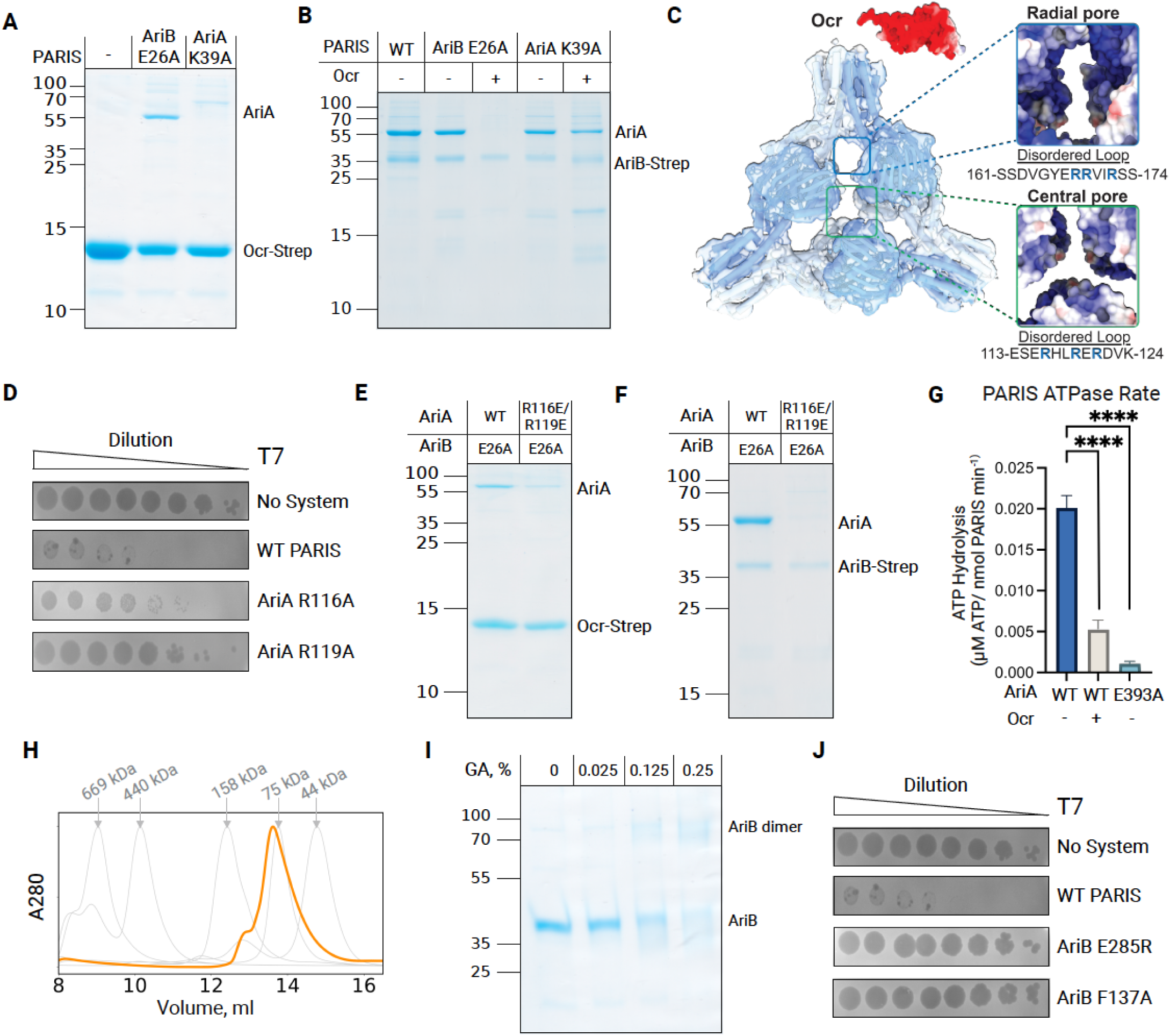
AriA interacts with the Ocr trigger in an ATPase-dependent manner, leading to the release of AriB from the PARIS complex. A) Ocr-Strep pulls down AriA, but not AriB. A mutation in the ATPase active site blocks binding of AriA to Ocr-Strep. B) Ocr releases AriB (E26A) from wild-type (WT) AriA (lane 3) but Ocr does not trigger AriB release from an ATPase-deficient variant of AriA (K39A) (lane5). C) A 4.5 Å-resolution reconstruction of AriA that was purified using Ocr-Strep as a bait. The structure reveals three radial pores that are symmetrically positioned around a central pore. The pores contain disordered loops with several positively charged residues. Ocr is shown as an electrostatic surface. D) Mutations in the central pore (i.e., R116A or R119A) reduce the efficiency of phage protection by several orders of magnitude. E) Ocr-Strep binds WT AriA and ejects AriB (E26A), while the charge swap mutations in the central pore of AriA (R116E/R119E) limit interaction with Ocr-Strep. F) AriB-Strep (E26A) pulls down WT AriA but not AriA (R116E/R119E). G) ATP and ADP were separated using thin layer chromatography and the rates of hydrolysis were measured by quantifying the accumulation of ADP (**Sup** Fig 6). The rates of ATP hydrolysis for PARIS, PARIS mixed with ten-fold excess Ocr, or PARIS with an ATPase mutation in AriA (E393A). The experiments were performed in triplicate. Statistical comparisons between groups were performed using a *post hoc* Dunnett’s test ****p<0.0001. H) Size exclusion chromatography of AriB-Strep (36 kDa) after Ocr-mediated release from AriA. The column was calibrated using molecular weight standards (gray lines). AriB elutes in single peak with an estimated molecular weight of 81 kDa (**Sup Fig 4A**). I) Glutaraldehyde (GA) cross-linking assay with activated AriB (E26A). J) Mutations predicted to block AriB dimerization (E285R and F137A) prevent PARIS-mediated defense (**Sup** Fig. 1G).

### AriA is an ABC ATPase that sequesters the AriB TOPRIM nuclease

While the homohexameric assembly of AriA was apparent in the early stages of reconstruction, conformational heterogeneity limited the resolution (**Sup Fig 3, Sup Movie 1**). To improve the resolution, we used local refinement to determine a 3.2 Å nominal resolution reconstruction of one asymmetric unit of the PARIS complex (AriA_2_:AriB_1_), into which we built an atomic model (**PDB ID: 8UX9, Sup Table 1**) (**Fig 2**). Each asymmetric unit of the PARIS complex is composed of two AriA molecules, which form a head to tail homodimer, and one AriB molecule that binds directly above the AriA dimer interface.

AriB is a metal-dependent TOPRIM nuclease and the predicted effector of the PARIS defense system^4,7,8^. The structure reveals a series of spatially conserved, negatively charged residues (E26, D30, D88, E90, and E122) that are characteristic of TOPRIM domains that bind two metal ions. The E26A mutation inactivates PARIS-mediated defense (**Fig 2B**)^4,7^. Glutamic acid 26 is just upstream of an N-terminal alpha helix (α2) on AriB, which includes two Arginine residues (R31 and R28) that form salt bridges with negatively charged residues on AriA that is positioned directly above the ATP binding sites (**Fig 2B, Sup Fig 2, Sup Movie 1)**. Mutations that interfere with the AriA:B interface (R31E, R28E) limit phage protection by PARIS (**Fig 2E**), which suggests that the protective activity of AriB requires an association with AriA.

The AriA ABC ATPase domain resembles that of Rad50 (RMSD 1.1 Å across 86 C-alpha atom pairs), a universally conserved protein involved in dsDNA break repair^19–21^. Structural comparison of AriA to Rad50 confirms that AriA contains the sequence motifs required for ATPase activity, along with two unique insertion sequences, IS1 and IS2 (**Fig 1B, Sup Fig 2**). Insertion sequence 2 contains a series of aromatic residues located on one face of helix α8 (F230, F234, F238, and Y304) that coordinate interactions between AriA homodimers, and explains how AriA homodimers assemble into a trimer of dimers (**Sup Fig 2**); a rare assembly state for this family of proteins^19^. Like other ABC ATPases, the head to tail assembly of the AriA homodimer results in two symmetrically positioned nucleotide binding domains (NBD) at the dimer interface (**Fig 2A)**. In the presence of a nonhydrolysable ATP analog (i.e., ATPγS), we observe density for the ligand in both nucleotide binding domains (NBDs), but in different coordination states (**Fig 2C-D**). Previous studies of ABC ATPases revealed that ATP hydrolysis occurs at the two NBDs in an alternating fashion^19,22^, which is consistent with the observed “open” and “closed” conformations of NBDs (**Fig 2C-D, Sup Movie 1**).

### AriA binds to the Ocr anti-restriction protein leading to AriB release and activation

The anti-restriction protein Ocr from phage T7 was previously identified as a trigger of the PARIS system^4,6^. To determine if AriA:B senses Ocr through a direct interaction, we co-expressed AriA with an active site mutant of AriB (E26A) and Ocr-Strep. AriA, but not AriB, co-purified with Ocr-Strep and the complex migrates according to the predicted mass of an AriA hexamer (**Fig 3A**, **Sup Fig 4A**). This is consistent with a direct interaction between AriA and Ocr, and release of AriB. A pull-down with AriB-Strep confirms that AriB dissociates from the complex in the presence of Ocr (**Fig 3B**).

**Figure 4:**
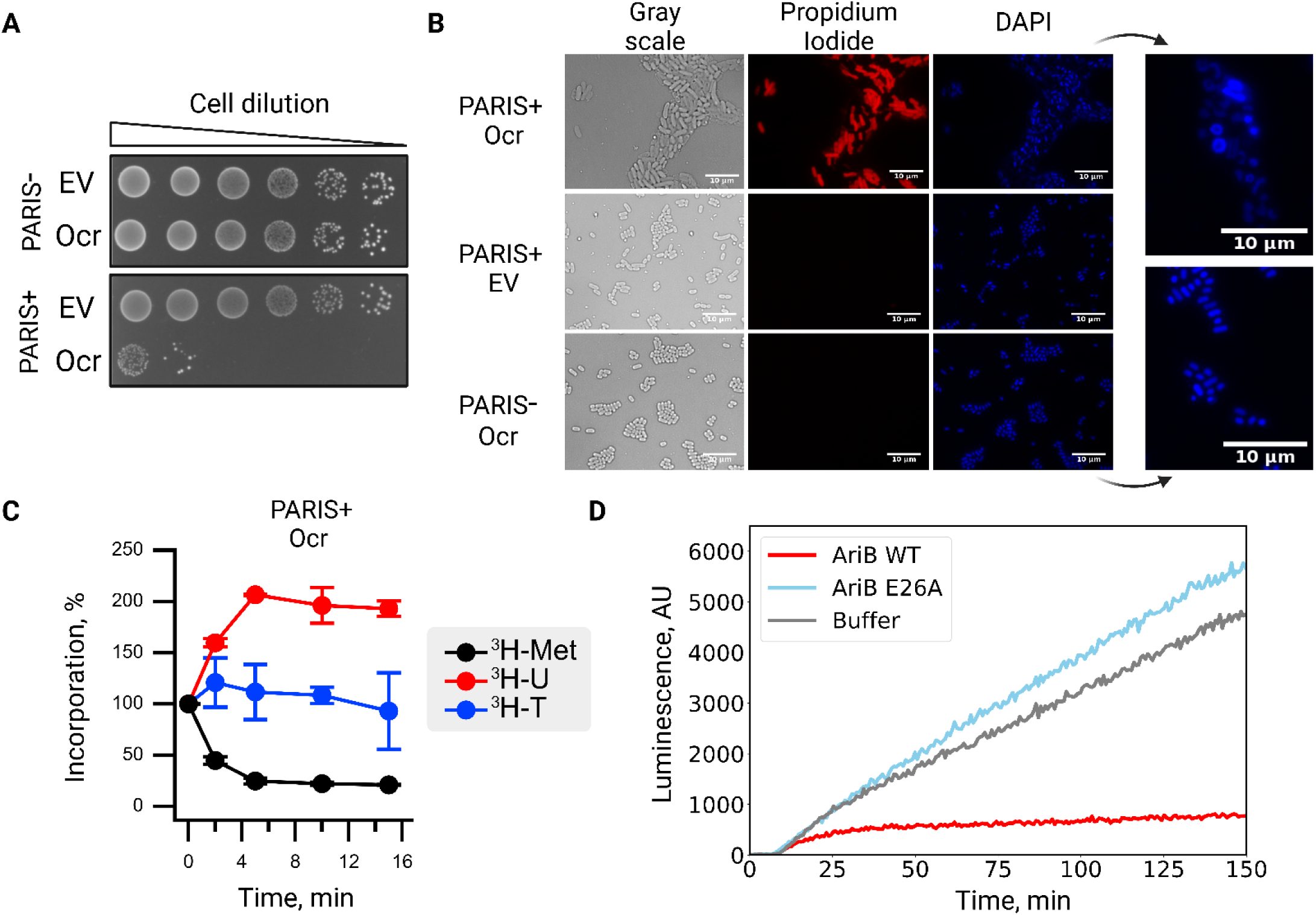
The activation of PARIS leads to cell death and inhibition of translation. A) Expression of Ocr from the pBAD vector in PARIS^+^ culture results in cell toxicity, while the empty vector (EV) has no impact. B) Ocr expression in PARIS^+^ cells induces cell death (propidium iodide staining, red) and DNA compactization (DAPI staining, blue). Arrows point to close-up images of the nucleoid structures from PARIS^+^ / Ocr (top) and PARIS^-^ / Ocr (bottom). Cells were imaged 1 hour after induction of Ocr expression. C) Metabolic labelling experiments reveal a drop in ^3^H methionine incorporation following induction of Ocr, indicative of translation inhibition upon PARIS activation. D) *In vitro* translation in the presence of activated AriB (WT) or active site mutant (E26A). Translation of firefly luciferase mRNA was monitored by accumulation of luminescent signal. In the presence of activated AriB, luciferase signal is reduced dramatically, further indicating that PARIS functions through a mechanism of translational inhibition.

To determine how Ocr triggers the release of AriB, we used the purified AriA:Ocr-Strep complex for structure determination. The 4.4 Å-resolution structure reveals a D3 symmetric, homohexamer of AriA with no AriB (**Fig 3С and Sup Fig 5**). At this resolution, the structure of AriA purified using Ocr-Strep is indistinguishable from the structure of AriA purified using AriB-Strep. We anticipated that the structure would include Ocr, since Ocr-Strep was used to pull down the AriA hexamer and the complex was stable during gel filtration. However, multiple data processing strategies failed to resolve Ocr, suggesting that Ocr has several binding sites on AriA and/or that the interaction is conformationally flexible.

**Figure 5:**
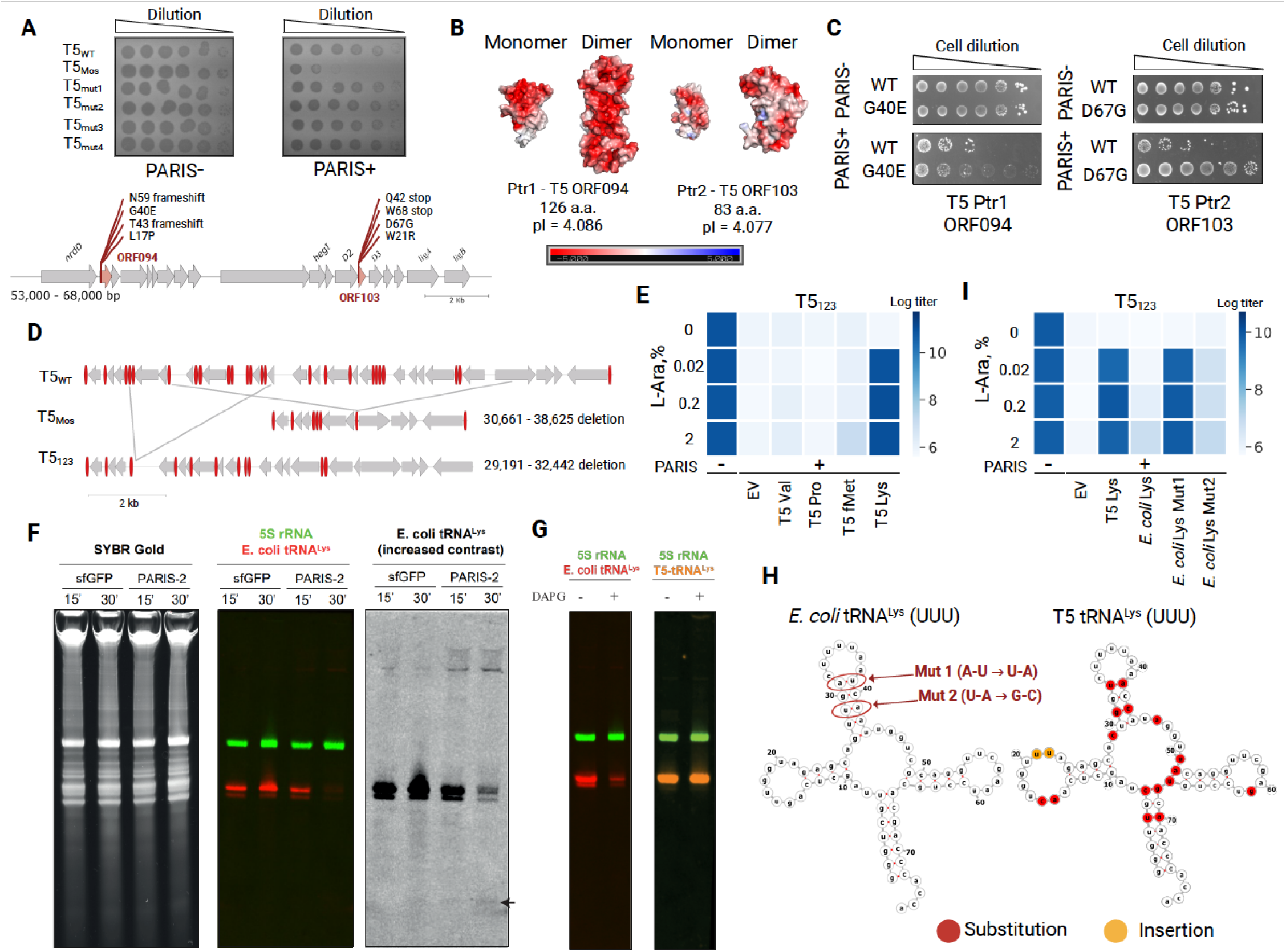
PARIS cleaves the *E. coli* tRNA^Lys^ and T5 bypasses immunity by encoding its own non-cleavable tRNA^Lys^. A) Serial dilution plaque assays conducted in PARIS +/- cells against T5_wt_, T5_Mos_ and T5_Mos_-derived PARIS escapers. T5_Mos_ phages insensitive to PARIS defense accumulate mutations in ORF094 and ORF103. B) AlphaFold2-predicted structures for T5-encoded PARIS triggers shown as monomers or dimers with their surfaces colored by electrostatic potential. C) Expression of Ptr1 (T5 ORF094) and Ptr2 (T5 ORF103), but not of the selected mutated variants, triggers PARIS toxicity. D) Comparison of T5_Mos_ and T5_123_ genomes with T5_WT_. T5_Mos_ is missing an 8 kb fragment while T5_123_ has a smaller 3 kb deletion. Both deletions encode genes for tRNA, shown in red. E) Overexpression of T5 tRNA^Lys^ in the PARIS^+^ cells rescues infection with T5_123_ phage that lacks tRNA^Lys^. F) Northern Blot performed on total RNA extracted from *E. coli* MG1655 carrying the PARIS system (pFR85) or a GFP control (pFR66), and *ocr* under the control of a P_PhlF_ promoter (pFD250). RNA was extracted 15 min or 30min after induction with DAPG. The 5S rRNA is marked with probe B811 (FAM) and the *E. coli* tRNA^Lys^ with probe B803 (IRDye700). A small black arrow shows a potential degradation product of the tRNA^Lys^. G) Northern Blot performed on total RNA extracted form *E. coli* carrying pFR85, pFD250 and the T5 tRNA^Lys^ under the control of a P_BAD_ promoter (pFD287). RNA was extracted after 30 min of incubation in LB with or without DAPG. The tRNA^Lys^ with probe B803 (IRDye700) and the T5 tRNA^Lys^ is marked with probe B806 (IRDye 800). H) Comparison of *E. coli* and T5 tRNA^Lys^ highlights substitution mutations near the anticodon stem loop that we hypothesize prevent targeting by AriB. I) *E. coli* tRNA^Lys^ carrying mutations in the anticodon stem loop, mimicking those of the phage T5, but not WT *E. coli* tRNA^Lys^, rescues infection of the T5_123_.

Since Ocr is a negatively charged DNA mimic^6^, we hypothesized that the binding site on AriA would be positively charged. An electrostatic surface of the AriA hexamer reveals a central pore, flanked by a series of three positively charged radial pores (**Fig 3C**). The radial pores contain two flexible loops (one from each of the adjacent AriA homodimers) with 14 unmodelled residues (161-174, SSDVGYERRVIRSS), while the central pore contains six loops (one from each AriA) that contain 12 unmodelled residues (113-124, ESERHLRERDVK). To determine the role of these loops in phage defense, we tested the effects of alanine substitutions and charge swap mutations on positively charged residues. Arginine to alanine substitutions (i.e., R116A and R119A) in the central pore result in a partial loss of phage protection (**Fig 3D**). Attempts to generate other mutations in the central (R116E, R119E) or radial pores (R168E, R168A, R172E, or R172A) of AriA consistently failed when we used a vector that also included wild-type AriB. We hypothesized that these mutations in AriA mimic trigger binding, which releases AriB, and results in toxicity. To test this hypothesis, we expressed the central pore mutant (R116E/R119E), with the catalytically inactive AriB (E26A), which is non-toxic. Pull down experiments performed with Ocr-Strep (**Fig 3E**) or AriB-Strep (**Fig 3F**) demonstrate that AriA (R116E/R119E) has limited interactions with both Ocr and AriB. Collectively, these results suggest that either the central, radial pore or both are involved in the interaction with the trigger.

Expression of AriB alone is not toxic to the cells, nor does it provide phage defense^4^. This suggests that the toxicity of AriB depends on its initial association with AriA. The ATPase activity of AriA is also required for phage defense^4^, which suggests that the ATPase of AriA is necessary for loading AriB onto AriA, or release of activated AriB. To differentiate between loading or release of AriB, we performed pull down assays using AriB-Strep and WT or an ATPase defective mutant of AriA (K39A). AriB-Strep associates with mutant and wild-type AriA with similar efficiency (**Fig 3B**), demonstrating that the ATPase mutant of AriA loads, but does not release AriB. We reasoned that the lack of AriB release in the presence of AriA K39A is due to a defect in trigger recognition. To confirm the role of AriA ATPase activity in trigger recognition, we conducted pull downs using Ocr-Strep and WT or K39A AriA. The AriA K39A mutant fails to co-purify with Ocr, indicating trigger recognition is ATPase dependent (**Fig 3A**). To determine if or how the trigger impacts the ATPase activity of AriA, we measured the rate of PARIS-mediated ATP hydrolysis in the presence or absence of Ocr and compared these results to an ATPase defective AriA mutant (**Fig 3G**). These results demonstrate that Ocr significantly (p=0.0001) reduces ATP turnover by AriA (**Fig 3G and Sup Fig 6**).

**Figure 6:**
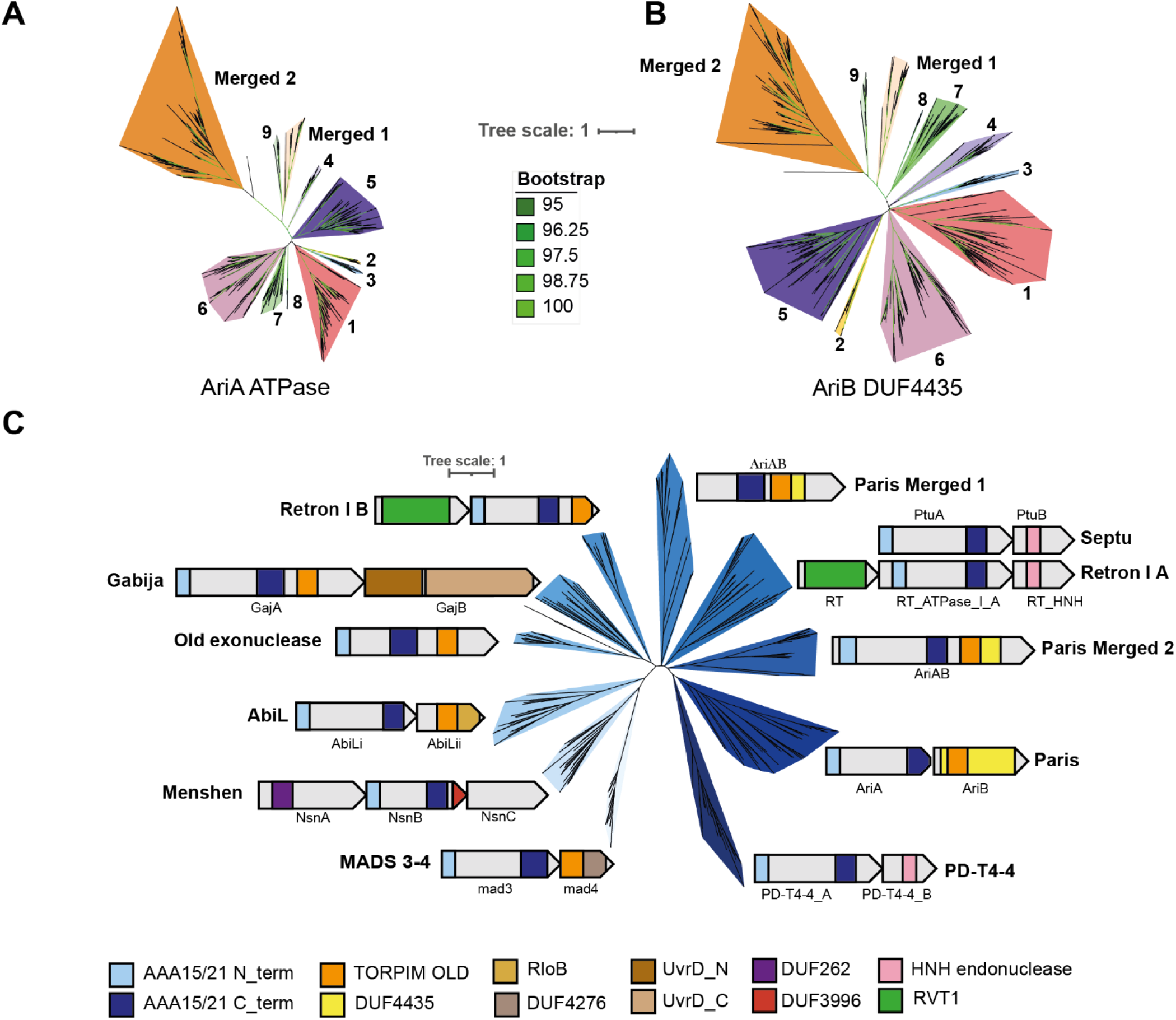
Phylogenies of PARIS and related ABC ATPase powered defense systems. A-B) Phylogenetic trees of AriA and AriB share similar branching patterns. Different colors represent different 11 subclades of PARIS including two clades in which *ariA* and *ariB* merged into a single *ariAB* gene (Merged 1 and Merged 2). C) Phylogenetic tree of the AAA15/21 ATPase containing defense proteins. Pfam accession: AAA15 (PF13175), AAA21 (PF13304), TOPRIM-Old (PF20469), DUF4435 (PF14491), RloB (PF13707), DUF4276 (PF14103), UvrD_N (PF00580), UvrD_C (PF13361), DUF262 (PF03235), DUF3996 (PF13161), HNH endonuclease (PF14279, PF10107), RVT1 (PF00078), were used for domain annotations.

TOPRIM-containing proteins are known to form homodimers^23,24^, but a predicted homodimer of AriB generated using AlphaFold2 clashes with AriA (**Sup Fig 1F)**. Thus, we propose that AriB forms an active homodimer after trigger-mediated release from AriA. The toxicity of PARIS upon activation makes purification of active AriB challenging. To overcome this problem, we mixed cell lysates expressing AriA:B-Strep and Ocr, and then recovered activated AriB using affinity chromatography (**Sup Fig 4B-C**). Size exclusion chromatography indicates that activated AriB forms a homodimer (**Fig 3H**, **Sup Fig 4A**), which was further confirmed by glutaraldehyde cross-linking experiments (**Fig 3I**). Mutations in the AriB dimerization interface (E285R and F137A), predicted from the AlphaFold2 model (**Fig 1B, Sup Fig 1G**), resulted in the loss of phage defense (**Fig 3J**), highlighting that AriB dimerization is essential for PARIS activity. Collectively, these data show that the Ocr trigger interacts with AriA in an ATPase dependent manner, leading to the release and dimerization of AriB.

### PARIS activation results in nucleoid compaction and translation inhibition

We then investigated the consequences of PARIS activation by Ocr. The expression of Ocr from a plasmid in *E. coli* is sufficient to cause cell death in the presence of PARIS (**Fig 4A**). Monitoring of PARIS^+^ cells under the microscope using live/dead staining with propidium iodide revealed the accumulation of cells with permeabilized membrane within 20 min of Ocr induction (**Fig 4B and Sup Movie 2**). The TOPRIM domain, likely responsible for AriB toxicity, is frequently associated with DNase or RNase activities^25^, but RNA or DNA preparations from AriB activated cells do not reveal evidence for indiscriminate nuclease activity (**Sup Fig 7A-C**). However, DAPI staining reveals that PARIS activation results in nucleoid compaction (**Fig 4B**), reminiscent of the translation inhibition phenotype^27,28^. To understand the nature of this compaction we performed chromosome conformation capture experiments (Hi-C). The results revealed a loss of chromosomal structures similar to that observed upon treatment with chloramphenicol, a known inhibitor of translation (**Sup Fig 8**).

To better understand the nature of PARIS-induced translational inhibition, we conducted metabolite labelling experiments and *in vitro* translation assays. Production of Ocr in PARIS^+^ cells resulted in rapid inhibition of ^3^H methionine incorporation, consistent with translation inhibition (**Fig 4C**), and in a moderate increase of ^3^H thymidine uptake. This increase is a known consequence of translation inhibition that can be explained by an enhanced transcription of rRNA genes^16^. The importance of activated AriB in translational arrest was further corroborated using a luciferase reporter in an *in vitro* translation assay. The presence of activated AriB in the reaction mixture interferes with the production of luciferase, while the active site mutant (i.e., AriB E26A) has no impact (**Fig 4D**). Collectively, the results demonstrate that PARIS activation results in translational inhibition.

### Phage T5 encodes two protein triggers and a suppressor of PARIS

The analysis of related phages has provided considerable insight into defense and anti-defense strategies^26^. While testing the activity of PARIS against phages of the T5 family, we noticed distinct plaquing phenotypes among related phages (**Fig 5A**). PARIS does not protect cells from T5_wt_ phage infection, but efficiently blocks infection by a variant of T5 from the Moscow phage collection (T5_Mos_). Consistent with translation arrest upon PARIS activation, infection of PARIS^+^ cells with T5_Mos_ at high MOI results in rapid growth inhibition, which occurs faster than the time it takes to lyse PARIS^-^ cells (**Sup Fig 9A**). Aside from a large deletion (*Δ30661-38625*), the T5_Mos_ genome is nearly identical to the T5 archetype (**Fig 5D**). We hypothesized that both phages contain a trigger that activates PARIS, but the T5 architype also contains a suppressor of PARIS that has been lost in T5_Mos_. To determine how T5_Mos_ triggers PARIS, we isolated and sequenced T5_Mos_ mutants that escape PARIS immunity. All PARIS-escape mutants carried mutations in two genes encoding highly acidic proteins of unknown function: ORF094 and ORF103 (**Fig 5A-B**). Expression of these proteins from a plasmid resulted in the activation of PARIS toxicity (**Fig 5C and Sup Fig 9**), confirming that T5 carries two novel PARIS triggers. Mutated variants encoded by escape mutants did not trigger PARIS. We named these proteins <u>P</u>ARIS <u>tr</u>igger 1 and 2 (Ptr1 - ORF094; Ptr2 - ORF 103)^27^. These results reveal that structurally diverse, negatively charged viral proteins can activate the PARIS system. The presence of *ptr1* and *ptr2* genes in the genome of T5_wt,_ which is not affected by PARIS, further indicates that this phage contains a suppressors of PARIS immunity.

### AriB cleaves the host tRNA^Lys^ and phage T5 encodes a non-cleavable tRNA^Lys^ substitute

The T5 genome encodes 23 tRNA, 16 of which have been lost in T5_Mos_. Given the inhibitory effect of activated AriB on translation, we hypothesized that phage-encoded tRNAs are responsible for overcoming PARIS toxicity. To narrow the search for a PARIS suppressor, we tested a collection of T5 deletion mutants for their sensitivity to PARIS^28^. Among the variants tested, T5_123_ carries the smallest deletion in the tRNA genes region (*Δ29191-32442*) and is still sensitive to PARIS (**Sup Fig 9B**). The DNA fragment missing from T5_123_ was divided into three segments, which were separately cloned on an expression plasmid and introduced in PARIS^+^ cells. A fragment encoding tRNA^Pro^, tRNA^fMet^, tRNA^Lys^ and tRNA^Val^, partially rescued the ability of phage T5_123_ to infect PARIS^+^ cells, while other fragments had no effect (**Sup Fig 9C**). Next, we cloned each of the tRNAs individually and tested their ability to rescue T5_123_ infection. Expression of the T5 tRNA^Lys^ completely restored T5_123_ infectivity (**Fig 5E**), demonstrating that this phage-encoded tRNA neutralizes PARIS-induced toxicity, and suggests that the host tRNA^Lys^ is targeted by AriB.

To confirm that the *E. coli* tRNA^Lys^ is indeed degraded upon PARIS activation, we performed a Northern Blot with total RNA extracted from PARIS^+^ cells 15 or 30 min after induction of the Ocr trigger. tRNA^Lys^ was degraded, while SYBR-Gold staining did not reveal degradation of other RNAs (**Fig 5F**). To determine how T5 tRNA^Lys^ suppresses PARIS immunity, we analyzed RNA extracted from PARIS activated cells expressing phage tRNA from a plasmid. Unlike *E. coli* tRNA^Lys^, the T5 tRNA^Lys^ remained intact (**Fig 5G).** The phage-encoded variant of tRNA^Lys^ contains mutations in the anticodon stem-loop (**Fig 5H**), which could be responsible for the lack of AriB recognition. To test this hypothesis, we introduced mutations from the T5 tRNA into the *E. coli* tRNA^Lys^ counterpart. The resulting chimeric tRNA rescued infection by phage T5_123_ in the presence of PARIS, confirming that AriB recognizes the tRNA anticodon loop (**Fig 5I**). Collectively, these results indicate that phage T5 encodes a non-cleavable variant of tRNA, compensating for the loss of the host tRNA depleted upon PARIS activation.

### Evolutionary history of the PARIS defense system

To inform our understanding of the diversity of PARIS systems and their relationship with other ABC ATPase powered defense systems, we conducted phylogenetic analyses. First, we generated trees for AriA and AriB, revealing a diverse set of systems that can be grouped into 11 distinct clades including two in which *ariA* and *ariB* merged into a single *ariAB* gene (**Fig 6A,B**). The *ariA* and *ariB* genes of a same system consistently fall in matching clades, showing they coevolve and are not frequently swapped between systems. A rooted version of the AriB tree shows that AriAB emerged once from clade 9 of two gene systems in a single event and was then split in two different clades (**Sup Fig. 10**). Interestingly, sequence alignments of AriA reveal that residues associated with the central and radial pores of the complex are poorly conserved (**Sup Fig 10**). This suggests that the signal transduction and effector parts of PARIS immunity are shared across homologs, while the central and radial pores have evolved to recognize different viral triggers.

PARIS was recently assigned to a family of OLD-like defense systems that share an ABC ATPase and a TOPRIM nuclease^12^. This analysis identified four classes, where the single gene systems such as P2 Old are categorized as class 1, Gabija are class 2, reverse-transcriptase containing systems are class 3, and PARIS systems are class 4. We built a tree of known anti-phage defense systems that carry an ABC ATPase of the AAA15/21 family (**Fig 6C**). Our analysis adds AbiL and MADS3-4 systems to the list of systems sharing an ABC ATPase and TOPRIM nuclease. It also reveals how this ABC ATPase can be found in association with a variety of other effector domains, revealing the evolutionary success of this domain as the likely sensor of viral infection in diverse immune systems.

## Discussion

Our results demonstrate how AriA forms propeller-shaped hexamer capable of binding three AriB monomers. AriA is an ATP-dependent sensor that detects viral immune suppressors (e.g., Ocr), which trigger the release of AriB. Released, AriB assembles into a homodimer that cleaves specific host tRNAs resulting in a rapid interruption of translation followed by a loss of membrane integrity. As opposed to prototypical toxin-antitoxin systems, AriB isn’t toxic when expressed alone. This suggests that AriA activates AriB through a structural modification that enables dimerization or an unidentified posttranslational modification. More work will be necessary to determine the mechanism of AriB release and activation.

Interestingly, we found that the PARIS system from *E. coli* B185 is not only triggered by phage T7’s Ocr protein, but also by Ptr1 and Ptr2 proteins from phage T5. Despite these proteins being small and negatively charged, they lack obvious sequence or structural similarities. Determining how PARIS detects such diverse proteins and whether different systems have distinct trigger sensitivities presents another intriguing avenue for future research.

Our work adds to a growing list of defense systems that trigger translational arrest upon infection. The PrrC and Retron I A (i.e., Ptu*AB* proteins) systems target tRNA degradation after detecting foreign proteins, while Cas13a targets tRNA degradation after detection of foreign RNA^29–31^. However, translational arrest isn’t limited to tRNA degradation, Mogila *et al* recently identified a CRISPR-Cas system that cleaves mRNA associated with the ribosome which arrests translation^32^, and the Lit (Late Inhibition of T4) protease degrades the translation elongation factor EF-Tu after detection of the major capsid protein of phage T4^33^.

We show that some phages circumvent PARIS-induced toxicity by expressing non-cleavable tRNA variants. Many tailed phages, such as those from *Demerecviridae* (T5-like phages), *Straboviridae* (T4-like phages) or *Ackermannviridae* families encode large arrays of tRNAs previously thought to be associated with translation optimization due to codon bias in the host and viral genome^34^. However, estimation of the abundance of host and phage tRNAs during infection, compared to the corresponding codon frequencies in produced mRNAs does not directly support this hypothesis^34^. A recent computational analysis suggests that phages may have evolved these variants in response to bacterial tRNA-targeting toxins, as evidenced by distinct anti-codon loop structures aimed at evading host nucleases^35^. This hypothesis is further supported by a parallel work demonstrating that phage T5 tRNA^Tyr^ provides protection against the PtuAB toxin of the Eco7 (Ec78) retron^31^.

Interestingly, this strategy of encoding tRNA clusters is also observed in some eukaryotic viruses, like those from *Herpesviridae, Mimiviridae* and *Phycodnaviridae* families^36^, possibly serving a similar function in evading host immunity. Strikingly, an anti-codon nuclease was also recently found to protect humans against Pox viruses, demonstrating that the inhibition of translation through the inactivation of specific tRNAs is an anti-viral strategy in each domain of life^37^.

## Methods

### Bacterial strains, Phages, and Plasmids

Phages, bacterial strains and plasmids used in the study are listed in Supplementary Table 2. Primers are listed in Supplementary Table 3. Infection with T5 phage was carried in LB supplemented with 1mM CaCl_2_. Unless otherwise indicated, 0.2% L-arabinose, 0.1mM IPTG, and 200 ng/mL aTc were used for the induction.

For structural studies, the coding sequences for AriA and AriB from *E. coli* B185^4^ were cloned under control of a T7 promoter into a pRSF-Duet vector with a C-terminal strep-tag II on AriB. For co-expression of PARIS with T7 Ocr trigger, the TOPRIM nuclease of AriB was inactivated (E26A) and the strep-tag II was removed by site-directed mutagenesis. The coding sequence for T7 Ocr protein was cloned into a pET-Duet vector with a C-terminal strep-tag II. For *in vitro* binding assays, the coding sequence for T7 Ocr protein was cloned into a pET-Duet vector with a N-terminal His6-TwinStrep-SUMO tag. All *in vivo* experiments were performed with pFR85, encoding PARIS from *E. coli* B185 with its native promoter and an inducible *Ptet* promoter upstream. For *in vivo* protein pull-downs, C-terminal AriB strep-tag II was introduced to the pFR85. Mutants of PARIS were constructed from pFR85 using the Gibson method^38^ or Q5 site-directed mutagenesis kit (NEB). The PARIS triggers Ptr1 (T5 ORF094) and Ptr2 (T5 ORF103) from T5 phage as well as T5 genome fragments, T5 tRNA^Lys^ and *E. coli* tRNA^Lys^ were cloned on the pBAD18 vector under control of *araBAD* promoter using the Gibson method. 20 bp upstream and downstream of tRNA genes were preserved during cloning. T5 nucleotide positions are provided according to the genome assembly AY543070.1. All constructions were verified using Sanger sequencing.

### Expression and purification of the PARIS complex

AriA and AriB-strep were co-expressed from a pRSF-Duet plasmid in BL21-AI cells (Invitrogen). Cultures grown at 37°C to 0.4-0.5 OD_600_ were induced with 0.1 mM IPTG and 0.1% L-arabinose. Following induction, the cultures were incubated overnight at 16°C. Cells were pelleted (3,000 × g, 10 min, at 4°C), resuspended in lysis buffer (25 mM Tris, pH=8.5, 150 mM NaCl, 1 mM EDTA) and lysed by sonication. Cellular debris was removed by centrifugation (10,000 × g, 25 min, at 4°C) and AriAB complex was purified by affinity chromatography on 5 mL streptactin resin (IBA) and eluted with 2.5 mM desthiobiotin (IBA) in lysis buffer. The eluted protein was concentrated (100k MWCO PES Spin-X UF concentrator, Corning) and further purified by size exclusion chromatography (Sup200 column, Cytiva) in lysis buffer with 2% glycerol. Fractions of interest were combined and concentrated to 5µM (100k MWCO PES concentrator, Pierce) and used immediately for vitrification on cryo-EM grids.

### *In vivo* protein pull-down

To determine protein-of-interest interacting partners *in vivo*, AriB-Strep expressed from pFR85 or Ocr-Strep expressed from pBAD were used as bait, and pull-downs were performed in *E. coli* BW25113. 3 L of cell culture was grown in LB media at 37°C in Thomson flasks with aerated lids (1.5 L per flask). Expression of proteins was induced with 200 ng/mL aTc or 0.2% L-arabinose at OD_600_ ∼ 0.1 and cells were harvested by centrifugation after reaching OD_600_ ∼ 0.8-1. Cells were processed as described above, and Strep-tagged proteins were purified on two stacked 5 mL StrepTrap HP (Cytiva) columns. Identity of protein bands after SDS-PAGE was determined by matrix-assisted laser desorption/ionization time-of-flight (MALDI-TOF) mass spectrometry. ∼1/3 of the Coomassie-stained band was cleaved from the gel and samples were prepared with Trypsin Gold (Promega) in accordance with manufacturer’s instructions. Mass spectra were obtained using the rapifleX system (Bruker).

For production of activated AriB, AriB-Strep from pFR85 and non-tagged Ocr from pBAD were expressed in separate cultures and cells were grown as described. Cell cultures were mixed, and cell walls were disrupted by sonication on ice (60% power, 10’’ pulse, 20’’ pause, 30-60 minutes) on Qsonica sonicator with 6 mm sonotrode, followed by StrepTrap HP purification of activated AriB-Strep. The molecular weight of protein complexes was determined after size exclusion chromatography performed on Superdex 200 Increase 10/300 column (GE Healthcare) calibrated with High Molecular Weight calibration kit (GE Healthcare).

### Glutaraldehyde cross-linking assay

To determine the oligomeric state of activated AriB in solution, we performed glutaraldehyde cross-linking assay. Concentrated AriB aliquots were mixed with glutaraldehyde (253857.1611; PanReac) to a final concentration of 0.025%, 0.125%, or 0.25% vol/vol and incubated for 30 min at room temperature. The reaction was quenched by the addition of one sample volume of 1 M Tris-HCl at pH 8.0 and a half sample volume of 5x concentrated SDS sample buffer. Samples were heated for 5 min at 60°C and separated by SDS-PAGE.

### *In vitro* protein pull-down

The T7 Ocr protein with an N-terminal His6-TwinStrep-SUMO was expressed from a pET-Duet plasmid in BL21(DE3) cells. Cultures grown at 37°C to 0.4-0.5 OD_600_ were induced with 0.5 mM IPTG. Following induction, the cultures were incubated overnight at 16°C. Cells were pelleted (3,000 × g, 10 min, at 4°C), resuspended in lysis buffer (25 mM Tris, pH=8.5, 150 mM NaCl, 1mM EDTA) and lysed by sonication. Cellular debris was removed by centrifugation (10,000 × g, 25 min, at 4°C) and HSS-Ocr was purified by affinity chromatography on 5 mL streptactin resin (IBA) and eluted with 2.5 mM desthiobiotin (IBA) in lysis buffer. The eluted protein was incubated with SUMO protease at 4°C overnight before washing over 5 mL Ni-NTA resin (25 mM Tris pH=7.5, 300 mM NaCl, 1 mM TCEP, 20 mM Imidazole) to remove SUMO-cleaved tag. Flow through protein was concentrated (10k MWCO PES Spin-X UF concentrator, Corning) and further purified by size exclusion chromatography (Sup6 10 300 column, Cytiva) in lysis buffer with 2% glycerol. Fractions of interest were combined and concentrated (10k MWCO PES Spin-X UF concentrator, Corning). Cleaved Ocr (3.8 uM) was incubated with WT PARIS (11.3 μM) at 37°C for 15 minutes before analysis via size exclusion chromatography (Sup6 10 300 column, Cytiva) in lysis buffer with 2% glycerol. Fractions of interest were combined, concentrated (10k MWCO PES Spin-X UF concentrator, Corning), and analyzed via SDS-PAGE (15% acrylamide, 150V for ∼1 hour).

### Cryo-EM sample preparation and data collection

All samples were subjected to vitrification on 1.2/1.3 Carbon Quantifoil^TM^ grids which were glow discharged using an easiGlow^TM^ (Pelco) glow discharge station (glow 45s hold 15s). After applying 3 µL of 5µM PARIS, grids were subjected to double-sided blotting using a Vitrobot Mk IV with a force of 5 for 4 seconds at 4° and 100% relative humidity. After plunge-freezing in liquid ethane, grids were clipped and stored in liquid nitrogen for subsequent analysis.

For the PARIS complex in the inactive state obtained from AriB pulldowns, cryo-EM grids were screened, and all data were collected using a Talos Arctica microscope (ThermoFisher) operating at 200kV equipped with a K3 direct electron detector (Gatan). SerialEM^39^ software version 3.8.6 was used to automate data collection. For final data collection, a total of 7,340 movies were collected in super resolution mode with a pixel size of 0.552Å/pixel and a total dose of 56.42 e/Å^2^ distributed over 50 subframes.

For *in vitro* Ocr pulldown data collection, samples were prepared for cryo-EM analysis as described above. After screening grids, 10,340 movies were collected using a Talos Arctica microscope (Thermofisher) operating at 200kV equipped with a K3 direct electron detector (Gatan). SerialEM software version 3.8.6 was used to automate data collection. Movies were recorded in super resolution mode with a pixel size of 0.552Å/pixel and a total dose of 56.31 electrons/Å^2^ distributed over 50 subframes.

### Cryo-EM Data Processing of Inactive PARIS

All data were processed in cryoSPARC v4.41. After patch motion correction and CTF estimation, micrographs with CTF fits worse that 8 Å were discarded, yielding 5,998 micrographs for particle picking. After processing a subset of 200 micrographs, the AriA_6_:AriB_3_ assembly of the PARIS complex was apparent. Using this low-resolution initial volume, Alphafold2 predicted structures of AriA and AriB were fit into the map and used to generate a 20 Å lowpass filtered map using the Molmap command in ChimeraX. This volume was imported to cryoSPARC and used to generate templates for particle picking. A total of 4,078,384 particles were extracted and after initial 2-D classification, 1,643,515 particles were kept for further processing. These particles were sorted using a three-class *ab initio* reconstruction which revealed two junk classes and a volume corresponding to the assembled PARIS complex that contained 940,782 particles. After a second round of 2-D classification, 934,763 particles remained were selected and subjected to another round of *ab initio* reconstruction and heterogeneous refinement. This produced a consensus volume containing 532,010 particles that contains conformational heterogeneity for two AriB subunits, while 1 asymmetric unit of the complex is well aligned. Masked local refinement was used to determine a 3.2 Å-resolution map of one asymmetric unit of the complex. By subjecting the consensus refinement with 532,010 particles was also subjected to multiple rounds of masked 3-D classification to isolate particles in the ‘cis’ and ‘trans’ arrangements. The fully assembled PARIS complex demonstrates flexibility based on 3DVA, which limited the attainable resolution of the fully assembled complex. The density map for both the local refinement and the full complex are available at EMDB-42719,43103, and 43104 and raw micrographs are available at EMPIAR-11832.

### Cryo-EM Data Processing of Ocr-Pulldown

All data were processed in cryoSPARC v4.41. Of the 10,340 collected micrographs, 9,399 were selected for further processing based on the CTF fit criteria described above. Using the blob picker with a particle diameter of 200 Å, particles were extracted and subjected to 2-D classification. In the 2-D classes, there were two distinct macromolecular complexes present, one of which was interpreted to be the AriA scaffold of the PARIS complex, while the other was interpreted to correspond to RNA polymerase (RNAP).

From seven selected classes corresponding to the AriA hexamer, 667,479 particles were selected for downstream refinement. From a 2-class ab initio reconstruction of these particles, 392,989 were kept. Iterative sorting by multi-class ab initio reconstruction and heterogenous refinements yielded a volume containing 62,732 particles. AriA subunits can be fit into this density, but the poor resolution makes this structure indistinguishable to the AriA scaffold identified in the intact PARIS complex. No density is observed for Ocr, which suggests multiple binding sites may be present on the PARIS complex, or that Ocr is attached to AriA in a flexible state. Density map available at EMDB-43105 and raw micrographs are available at EMPIAR-11833.

From 6 selected classes corresponding to RNAP, 548,741 particles were subjected to a 2-class ab initio reconstruction. After removing a junk class, 371,555 particles remained. Further sorting using iterative ab initio reconstructions and heterogenous refinements isolated a stack of 190,177 particles. Non uniform refinement of these particles reveals a density map that closely resembles the structure of RNAP but suffers from severe anisotropy, preventing identification of bound Ocr.

### Model Building and Refinement

To determine the structure of the asymmetric unit of the PARIS complex, density halfmaps were obtained from cryoSPARC local refinements before being provided as inputs to DeepEMhancer for map sharpening. Alphafold predicted structures of AriA and AriB were then docked into the map density and map to model fit was improved using ISOLDE’s molecular dynamics simulation environment in ChimeraX. After initial equilibration, the PDB files were saved and used as inputs for realspace refinement using PHENIX. After initial refinement, problematic areas were manually inspected and modified using COOT and sidechains were removed from regions with poorer than 4 Å-resolution. After addressing clashes and non-rotameric sidechain orientations, iterative refinements were carried out in Phenix until Clashscores, Ramachandran and sidechain outliers failed to improve. The final model and their corresponding density map was submitted to the PDB for deposition (PDB ID: 8UX9, EMDB-42719,

To generate the biological assemblies for the PARIS complex atomic model, halfmaps from C3 symmetric, and nonuniform refinements of particles corresponding to the ‘cis’ and ‘trans’ orientations were obtained from cryoSPARC and sharpened in DeepEMhancer. After fitting the three copies of the asymmetric unit into the ‘cis’ and ‘trans’ density maps using the ChimeraX fitmap command, the find NCS tool in Phenix was then used to calculate the appropriate symmetry operators for the assembled forms of the complex. These symmetry operators were applied to the Asymmetric unit to generate biological assembly 1 and 2 using the Apply NCS tool in Phenix. Maps for the ‘cis’ and ‘trans’ orientations of the PARIS complex are available at EMDB-43104 and EMDB-43105, respectively.

### ATPase Assays

ATPase activity was measured using 6uM Ocr, 100 nM PARIS and 128 µM ATP supplemented with 10 nM [α-32 P]-ATP (PerkinElmer). The reaction buffer includes 20 mM Tris-HCl pH 8.0, 150 mM Sodium Chloride, 1 mM DTT, and 5 mM magnesium chloride. The reactions incubated at 37°C and 10 uL aliquots were quenched with Phenol at 1, 2, 4, 8,16, and 32 minutes. For the ‘trigger’ only reactions, 100 nM Ocr was mixed with 128 µM ATP supplemented with 10 nM [α- 32 P]-ATP (PerkinElmer) in the reaction buffer. Reactions were incubated for 32 minutes at 37°C and quenched with phenol. The ATP and ADP makers were created using T4 polynucleotide kinase (NEB). T4 PNK mixed with 128 µM ATP supplemented with 10 nM [α-32 P]-ATP, and 1 mM DNA oligo in T4 polynucleotide kinase buffer (NEB). This mixture was then incubated at 37°C for 1 hour. All reaction products were phenol-chloroform extracted and resolved on silica TLC plates (Millipore). Reaction products were spotted 2.5 cm above the bottom of the TLC plate. The plate was placed in a TLC developing chamber filled to ∼1.5 cm with developing solvent (0.2 M ammonium bicarbonate pH 9.3, 70% ethanol, and 30% water) and covered with aluminum foil for 4 hrs at room temperature. The TLC plate was exposed to a phosphor screen, imaged with Typhoon phosphor imager and quantified with the ImageQuant software package (Cytiva).

### Efficiency of Plaquing Assay

The activity of PARIS mutants was measured in comparison to the wild-type system by performing efficiency of plaquing (EOP) assays with phage T7. *E. coli* K-12 MG1655 carrying each of the AriA or AriB mutants, the control plasmid pFR66 (sfGFP), or the wild-type PARIS system (pFR85) were grown overnight in LB + kanamycin 50 µg/ml. Bacterial lawns were prepared by mixing 100 µl of a stationary culture with 5 ml of LB + 0.5% agar, and the mixture was poured on Petri dish of LB + kanamycin 50 µg/ml. Ten-fold serial dilutions of high titer stock of T7 phage were spotted on each plate and incubated 5h at 37°C. EOPs with T5 and T5-like phages were performed with *E. coli* BW25113 as described above, except that plates were incubated at 37°C overnight, and top agar was supplemented with 1mM CaCl_2_. Induction of tRNA genes or phage T5 fragments was achieved by supplementing top agar with indicated amount of L-arabinose. Plaque assays were performed in at least two independent replicates.

### Liquid culture phage infection

To monitor dynamics of T5_Mos_ phage infection of the PARIS^+^ (pFR85) and PARIS^-^ (pFR66) culture in liquid, we used EnSpire Multimode Plate Reader (Perkin Elmer). Overnight bacterial cultures were diluted 100-fold in LB medium with appropriate antibiotics and grown at 37°C in 10 mL of LB supplemented with 1mM CaCl_2_. At OD_600_ = 0.6, 200 μl aliquots were transferred to 96-well plates and infected at indicated Multiplicity of Infection (MOI). Optical density was monitored for 10 hours. All experiments were performed in three biological replicates.

### Toxicity assay

PARIS toxicity in the presence of T7 Ocr or Ptr1 and Ptr2 was measured in a spot-test assay. PARIS^+^ (pFR85) and PARIS^-^ (pFR66) cultures carrying pBAD vector encoding indicated trigger were grown overnight in 10 ml LB at 37°C. Stationary cultures were diluted to OD=0.6 and plated on LB or minimal media (M9 with 5% v/v LB) agar plates supplemented with 0.2% L-arabinose by serial 10-fold dilution. Control plates without induction contained 0.2% glucose to prevent leakage of *araBAD* promoter.

### Fluorescence microscopy

PARIS^+^ (pFR85) and PARIS^-^ (pFR66) carrying pBAD Ocr were diluted 1:100 and cultivated in LB supplemented with appropriate antibiotics at 37°C. When OD_600_ reached 0.4, an aliquot was mixed with DAPI (Invitrogen) at 1 mg/mL final concentration and propidium iodide (PI, Invitrogen) at 1 μg/mL final concentration and incubated at room temperature in the dark for 5 minutes. LB + 1.5% agarose slabs (∼0.2 mm thick) were prepared on a 75 x 25 mm microscopy slide (Fisher Scientific). Agarose slabs contained staining dyes at the same concentration and optionally were supplemented with 0.2% L-arabinose, to induce Ocr expression. ∼1 μl of stained cells was placed on agarose slab and after 1-min drying the slab was covered by a small 22 × 22 mm coverslip (Fisher Scientific). Imaging was performed on a Nikon Eclipse Ti-E inverted epifluorescence microscope. Each field of view was imaged in transmitted light channel (200 ms exposure) and in DAPI (filterset Semrock DAPI-50LP-A, 200 ms exposure) and in PI (filterset TxRed-4040C, 200 ms exposure) channels.

### Extraction of Total DNA

Total DNA was extracted at different time points (15, 30, 60 and 120 min) with or without DAPG to induce the expression of Ocr in the presence of PARIS. DNA was extracted using the Wizard® genomic DNA purification kit (Promega) according to the manufacturer’s instructions and treated with RNase at 0.1 mg/ml final. Each sample of total DNA was loaded on 1% agarose 1X TAE gel.

### TUNEL assay

TUNEL assays was performed to estimate *in vivo* accumulation of dsDNA breaks, according to methods described previously^40^. PARIS^+^ (pFR85) culture carrying pBAD Ocr vector was grown to OD_600_=0.3, followed by Ocr induction with 0.2% L-arabinose for 2 hours. As a positive control, cells were treated with 0.1% H_2_O_2_ to induce accumulation of DNA breaks. 2 ml of cells were harvested by centrifugation and treated according to the standard protocol (TUNEL Assay Kit – FITC, ab66108, Abcam). Measurement of FITC fluorescence was performed with CytoFLEX cytometer (Beckman Coulter) in the FITC channel. 100,000 events were collected for each sample in three biological replicates. Data was analyzed and visualized in FlowJo v10.

### Hi-C Procedure and Sequencing

*E. coli* MG1655 carrying plasmids pFR85 (P_tet_ ariAB) and pFD250 (P_PhlF_ ocr), or the pFR66 (sfGFP) + pFD245 (PhlF sfGFP) control plasmids were diluted 1:100 from an overnight culture in LB + kanamycin 50 µg/ml + chloramphenicol 20 µg/ml and grown until OD_600_ = 0.3. DAPG 50 µM was then added, followed by incubation for 15 and 30min. To compare the effect of PARIS activation with that of treatment with chloramphenicol, *E. coli* MG1655 harboring pFR66 were diluted 1:100 from an overnight culture in LB + kanamycin 50 µg/ml until OD_600_ = 0.3, followed by addition of chloramphenicol 20 µg/ml and incubation for 15 and 30min. Cell fixation was performed with 4% formaldehyde (Sigma-Aldrich, Cat. F8775) as described in Cockram et al. (2021)^41^. Quenching of formaldehyde with 300 mM glycine was performed at 4°C for 20 min. Hi-C experiments were performed with the Arima kit. Samples were sonicated using Covaris (DNA 300bp).

Preparation of the samples for paired-end sequencing was performed using Invitrogen TM Colibri TM PS DNA Library Prep Kit for Illumina according to manufacturer instructions. The detailed protocol is available in Cockram et al. (2021)^41^. Reads were aligned with bowtie2 v2.4.4 and Hi-C contact maps were generated using hicstuff v3.0.3 (https://github.com/koszullab/hicstuff) with default parameters and using HpaII enzyme to digest. Contacts were filtered as described in^42^, and PCR duplicates (defined as paired reads mapping at exactly the same position) were discarded. Matrices were binned 4kb. Balanced normalizations were performed using ICE algorithm^43^. For all comparative analyses, matrices were downsampled to the same number of contacts. The Hi-C signal was computed as the contacts between adjacent 5kb bins as described in Lioy *et al*. (2018)^44^. In order to compare this signal with other genomics tracks, we binned it at the desired resolution.

### Metabolic labeling

Metabolic Labeling with 3H-methionine, 3H-uridine and 3H-thymidine Metabolic labelling experiments were performed as described previously^45^, with minor modifications. *E. coli* BW25113 PARIS^+^ (pFR85) strain was transformed with pBAD plasmid carrying Ocr trigger (for L-arabinose-inducible expression). Transformed cells were initially plated on LB plates supplemented with 100 μg/mL ampicillin, 25 μg/mL kanamycin, and 0.2% glucose. Using individual *E. coli* colonies for inoculation, 2 mL liquid cultures were prepared in defined Neidhardt MOPS minimal media (supplemented with 100 μg/mL ampicillin, 25 μg/mL kanamycin, 0.1% casamino acids, and 1% glucose, and grown overnight at 37°C with shaking. Subsequently, experimental 20 mL cultures were prepared in 125 mL conical flasks in MOPS medium, supplemented with 0.5% glycerol, 100 μg/mL ampicillin, 25 μg/mL kanamycin, as well as a set of 19 amino acids (lacking methionine), each at a final concentration of 25 μg/mL. These cultures were inoculated at OD_600_ of 0.05 and grown at 37°C with shaking until the OD_600_ reached 0.3. At this point, 1 mL aliquots (designated as the pre-induction zero time-point) were transferred to 1.5 mL Eppendorf tubes containing 10 μL of pre-incubated at 37°C respective radioisotope (3H methionine [0.77 μCi, Perkin Elmer], 3H uridine [0.1 μCi, Perkin Elmer] or 3H thymidine [0.32 μCi, Perkin Elmer]). Concurrently, Ocr expression in the remaining 19 mL culture was induced by adding L-arabinose to a final concentration of 0.2%. Throughout the Ocr induction time course, 1 mL aliquots were taken from the 20 mL culture and transferred to 1.5 mL Eppendorf tubes containing 10 μL of the appropriate radioisotope. Radioisotope incorporation was halted after 8 minutes of incubation at 37°C by adding 200 μL of ice-cold 50% trichloroacetic acid (TCA) to the 1 mL cultures. Additionally, 1 mL aliquots were sampled for OD_600_ measurements during induction time course. The resultant 1.2 mL TCA-stopped culture samples were loaded onto GF/C filters (Whatman) prewashed with 5% TCA and unincorporated label was removed by washing the filter twice with 5 mL of ice-cold TCA followed by a 5 mL wash with 95% EtOH (twice). The filters were placed in scintillation vials, dried for two hours at room temperature, followed by the addition of5mLEcoLite™-scintillation cocktail (MP Biomedicals). After shaking for 15 minutes radioactivity was quantified using automatic TDCR Liquid Scintillation Counter (HIDEX). Isotope incorporation was quantified by normalizing radioactivity counts (CPM) to OD_600_, with the pre-induction zero time-point serving as the reference (set to 100%). All experiments were performed as biological triplicates using three independent liquid starter cultures inoculated with different colonies.

### *In vitro* translation

*In vitro* translation was monitored by the production of luciferase signal in a PURExpress *in vitro* protein synthesis kit (NEB), using firefly luciferase mRNA as an input. pT7-Fluc, encoding firefly luciferase (*fluc*) gene under control of T7 promoter, was used as a template for T7 *in vitro* transcription with MEGAscript kit (Thermo Scientific). mRNA was treated with DNase I (Thermo Scientific) and purified with Monarch PCR & DNA cleanup kit (NEB). PURExpress *in vitro* translation reaction was assembled in RNase-free tubes according to manufacturer’s instructions, with minor modifications. Reaction mixtures were adjusted to a total volume of 5 μl (2 μl solution A; 1,5 μl solution B; 0,3 μl RiboLock (40 U/μl, Thermo Scientific); 0,2 μl D-Luciferin (10 mM, Sigma)), supplemented with 0.5 μl of activated AriB or AriB (E26A) (60 μg/ml final concentration) or the same volume of buffer A (50 mM NaCl; 1mM EDTA; 100 mM Tris-HCl; 5mM β-ME; pH 8) as a control. Reaction was started by the addition of 0.5 μl of purified *fluc* mRNA (500 ng), immediately placed in 384-well white plate and covered with optically clear film. Accumulation of luminescent signal was measured by EnSpire Multimode Plate Reader (PerkinElmer) for 3h at 37°C. AriB concentration was determined with Qubit Protein Broad Range Assay Kit (Invitrogen).

### Isolation and sequencing of T5 escaper mutants

To select PARIS escape mutants, phage T5_Mos_ was continuously incubated with PARIS^+^ (pFR85) culture for 3 days. In short, after reaching an OD_600_ = 0.6, cells were infected with T5_Mos_ at MOI∼0.1, and the culture was incubated overnight at 37°C. Phage was collected the next day and 1 ml of lysate was used to initiate the next round of infection with fresh PARIS^+^ (pFR85) culture. Escaper mutants were re-purified from the single plaques obtained on PARIS^+^ (pFR85) culture and produced on PARIS^-^ (pFR66) culture. 8 mL of high-titer lysate (∼10^10^ pfu/ml) was PEG-precipitated, and phage genomic DNA was purified as described previously^46^. DNA libraries were prepared according to a standard protocol and sequenced on MiniSeq platform (Illumina) with paired-end 150 cycles (75 + 75). Genome assemblies were performed with SPAdes implemented in Unicycler^47^. To identify mutations, genomes of T5_Mos_ escapers were aligned with the sequenced genomes of T5_Mos_ and T5_wt_ from initial stocks. T5 strains reported by Glukhov et. al.^28^ were sequenced on DNBSeq-G400 platform (BGI) to validate boundaries of deletions.

### Extraction of Total RNA

Total RNA was isolated from *E. coli* strain MG1655 harboring pFR85 (P_tet_ ariAB) or a control plasmid pFR66 (P_tet_ sfGFP). Cells also carried pFD250 (P_PhlF_ ocr) with Ocr under the control of an inducible DAPG P_PhlF_ promoter (**Fig 5F and Sup Fig 7B**), as well as plasmid pFD287 carrying the tRNA^Lys^ of phage T5 under the control of P_araBAD_ promoter inducible by arabinose (**Fig 5G**).

Cells were diluted 1:100 in LB with the appropriate antibiotics from overnight cultures and grown with agitation at 37°C. When cultures reached an OD_600_ of 0.2, aTc was added to a final concentration of 0.5 µg/ml to induce the expression of the PARIS system. For the experiment shown in Fig 5G, arabinose was added to a final concentration of 0,2% to induce the T5-tRNA^Lys^ from plasmid pFD287. Growth continued until 0.4 and the Ocr trigger was induced with DAPG added to a final concentration of 50 µM, followed by incubation for an additional 15 and 30mn post induction.

Cells were centrifuged at 4,000 g for 10 min, resuspended with 0.2 ml of lysozyme buffer (20 mM Tris-HCl, pH8.0, 2 mM EDTA, 1% Triton X-100 and 20 mg/ml lysozyme) and lysed for 30 min at 37°C. One milliliter of TRIzol reagent (Zymo Research) was added to the samples and incubated for 5 min at room temperature (RT). Lysates were extracted with 0.2 ml of cold chloroform and centrifuged at 12,000 g for 15 min at 4°C. The aqueous phase of the sample was precipitated with 0.5 ml of 100% cold isopropanol, incubated 10 min at RT and centrifuged at 12,000 g for 10 min at 4°C. RNA pellets were washed with 1 ml of 75% ethanol, dried at RT and dissolved in 50 µl of nuclease free water. Total RNA concentration was measured with a nanodrop spectrophotometer and then the samples were resolved in a 7 M Urea 10% acrylamide gel before visualization with SYBR-Gold.

### Northern Blot

RNAs were run for 1 hr at 180V in a 7M Urea 10% acrylamide gel and then transferred to a nylon membrane (Invitrogen) using XCell SureLock Mini-Cell system with the XCell II Blot Module (Invitrogen). The membrane was cross-linked 5 min with UV. After prehybridization with 30 ml of solution containing 6 X SSC, 0.5% SDS and 0.05 % casein for 1h at 45°C, the membrane was hybridized with 20 µM FAM or IR dye-labeled oligonucleotide probes (**Sup Table 2**) in 10 ml of prehybridization solution at 45°C overnight. Finally, the membrane was washed twice with 2 X SSC, 0,1% SDS for 10 min at RT and twice with 0.2 X SSC, 0.1% SDS for 10 min at 60°C and images captured using a ChemiDoc MP Imaging system (BIO-RAD).

### Defense system detection

Defense system detection was performed using DefenseFinder v1.1.1^48^ and DefenseFinder models v1.2.3 on all RefSeq complete genomes from July 2022.

### Domain annotation

Defense system domain annotation was done using HHpred^49^ webserver with the PFAM^50^ database and standard setting. For PARIS, trigger binding domain and Coiled-coil domain were annotated using the structure information and ESMfold^51^ structure prediction for AriAB.

### AriA and AriB tree

PARIS systems detected by DefenseFinder v1.1.1 on RefSeq complete genomes from July 2022 were used to build phylogenetic trees. Only hits to AriA and AriB with a coverage of more than 75% were kept to avoid pseudogenes. Hits were then clustered using Mmseqs2^52^ v13.45111 on AriA and AriAB with an identity threshold of 80% and coverage threshold of 90%. A single representative of each cluster was then used to build the AriA tree, and the cognate AriB was used to build the AriB tree. The AriA tree was built using the AAA15 ATPase domain only and the AriB tree using the DUF4435 domain only. Domains were detected by HMMsearch (from HMMER 3.3.2) on the *E. coli* B185 PARIS system and a first alignment using Mafft v7.505^53^ (default parameters) was used to extract the corresponding sequences in other PARIS variants. The final alignment used to make the tree was done using MUSCLE v.5.1.linux64^54^ using the model super5 and trim using clipkit v1.3.0 with option seq-gap. The DUF4435 tree using M5 Ribonuclease as outgroup was done using selected representative hits from different clades and aligned using muscle v5.1.linux64 with model super5. All trees were built using Iqtree v2.0.6^55^ using ModelFinder and Ultrafast Bootstrap 1000.

### Defense system ATPase AAA15/21 tree

ATPase AAA15/21 defense system containing detection was done using hmmsearch from HMMER 3.3.2 and Pfam Hidden Markov Model 33.1 on defense systems previously detected. Random selection of 20 proteins from the detection results with a coverage of more than 75% were selected in addition to experimentally validated homologs to build the tree. The ATPase alignment was done using Mafft v7.505 and the tree was built using Iqtree v2.0.6 using ModelFinder and Ultrafast Bootstrap 1000.

## Acknowledgements

Thanks to members of the B.W. laboratory for feedback and discussions, Coltran Hophan-Nichols and the cyber security team at Montana State Univeristy for computational support, Dr. Heather Callaway for helpful suggestions for data processing, Dr. Martin Lawrence and Colin Gauvin for maintenance and operation of the Cryo-EM Core Facility at Montana State University. The cryo-EM Facility at Montana State University is supported by NSF 1828765 and the M.J. Murdock Charitable Trust. Research in the Wiedenheft lab is supported by the National Institutes of Health (R35GM134867), the M.J. Murdock Charitable Trust, a young investigator award from Amgen, and the Montana State University Agricultural Experimental Station (USDA NIFA). N.B. is support by an F31 from the NIH (GM153146). A.S-F. is supported by NIH (K99GM147842) and the Burroughs Wellcome Fund (G-1021106.01). A.B.G. is supported by Montana State University’s Undergraduate Scholars Program, and by the NIH NIGMS IDeA program (P20GM103474). Skolkovo Institute of Science and Technology team was supported by the Ministry of Science and Higher Education grant (075-10-2021-114) and RSF grant (22-14-00004). We thank Anatoly Glukhov for sharing a collection of T5 deletion variants, Tinashe Prince Maviza for the help with *in vitro* translation assays, and Alina Demkina for the help with BGI sequencing. Illumina sequencing has been performed at the Skoltech Genomics Core Facility. D.B and F.D are supported by the European Research Council [677823]; European Research Council [101044479]; Agence Nationale de la Recherche [ANR-10-LABX-62-IBEID]. Molecular graphics and analyses performed with UCSF ChimeraX, developed by the Resource for Biocomputing, Visualization, and Informatics at the University of California, San Francisco, with support from National Institutes of Health R01-GM129325 and the Office of Cyber Infrastructure and Computational Biology, National Institute of Allergy and Infectious Diseases. Funders had no role in the conceptualization, designing, data collection, analysis, decision to publish, or preparation of the manuscript.

## Data availability

All information is available upon request to the lead contacts. Sequencing data have been deposited in NCBI database under BioProject id PRJNA1040033.

EM maps of PARIS in various assembly states and the atomic model for the asymmetric unit of the complex were deposited to the Electron Microscopy Data Bank (EMDB) and Protein Databank (PDB) databases. Accession codes can be found in Supplementary Table 1 of the manuscript. The PDB code for the experimentally determined structure of the PARIS asymmetric unit is 8UX9. EMDB Accession codes EMD-42719, EMD-43103, EMD-43104, and EMD-43105. Raw micrographs are available at EMPIAR-11832 and EMPIAR-18833. All constructs (wild-type and mutants) used in this study can be obtained on request to the lead contacts.

