## Supplementary Figures 1-9 for "Viral proteins activate PARIS-mediated tRNA degradation and viral tRNAs rescue infection"

**Supplemental Figure 1: Alphafold2 structural prediction of AriA and AriB.**

Alphafold2-predicted structures of AriA and AriB oligomers colored by the predicted local distance difference test (pLDDT) score (0-100) with associated scale bars. Values greater than 90 indicate high confidence and values below 50 indicate low confidence. Predicted Aligned Error plots (PAE) are also shown for each structural prediction. A) Alphafold2-predicted structure of the AriA homodimer. Confidence is high (PLDDT >90, green) for the ATPase domain, and low for the insertion sequences (PLDDT<70, teal). B) Alphafold2-predicted structure of the AriB homodimer returns a high confidence model with more >98% of the residues above a PLDDT of 90. C) Alphafold2-predicted structure of the AriA-B heterodimer and associated PAE plot. D) Alphafold2 returned unreliable models when attempting to predict structures for higher ordered assemblies of AriA and AriB as demonstrated by the PLDDT plot below 50 for most of the molecule. E) Clashing (red) between AriBs, prevents the assembly of two AriB subunits on a single AriA homodimer. Clashing is defined by atoms with overlap greater than 0.6 Å. F) A predicted homodimer of AriB cannot associate with a homodimer of AriA without clashing (red). Atoms with an overlap greater than 0.6 Å overlap are highlighted with red disks signifying the clash. G) Predicted structure of the AriB dimer shown in panel B with residues involved in dimerization indicated.

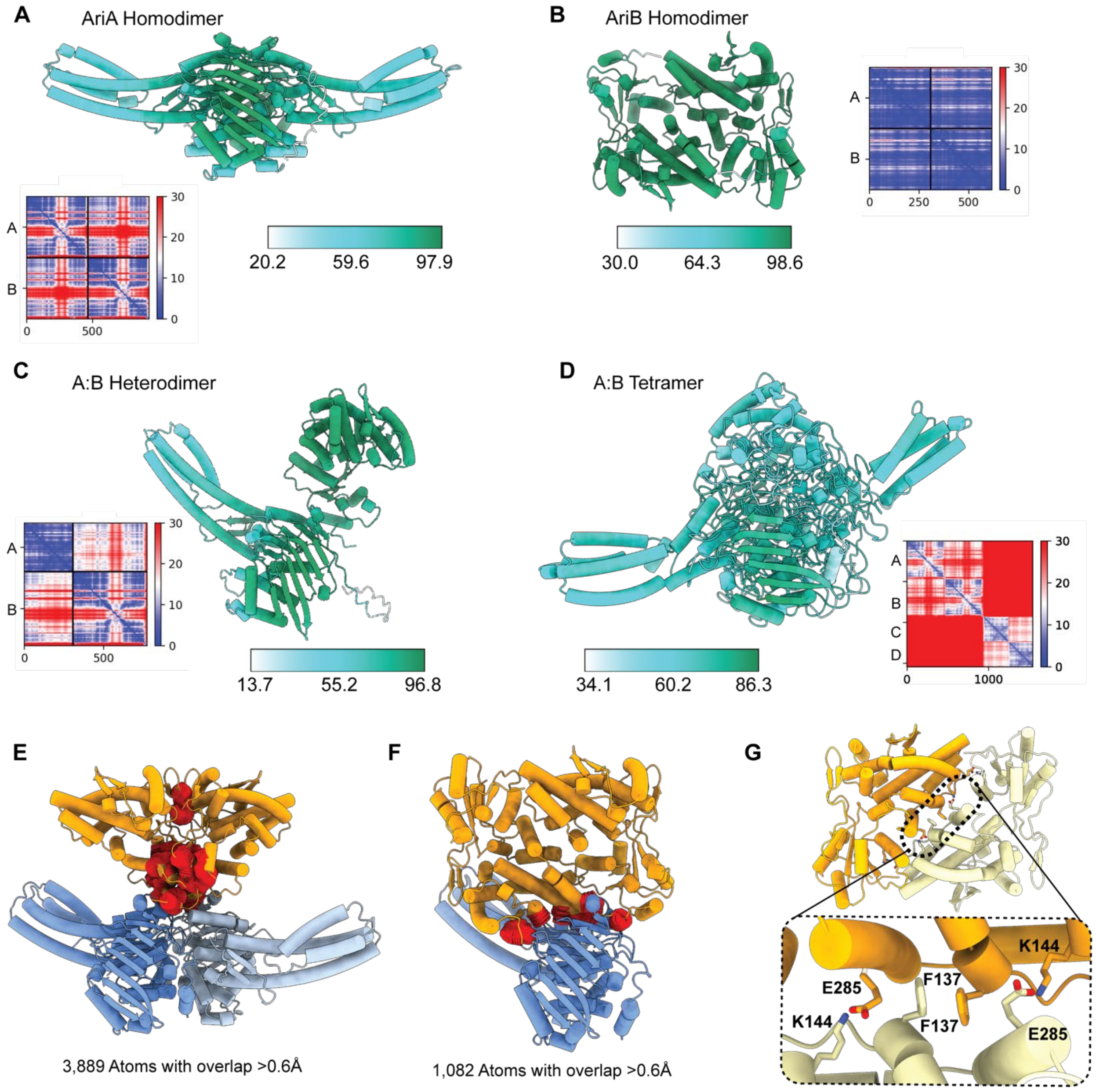

**Supplemental Figure 2: Structural comparison of PARIS homologs**

A) Relative domain organization of phage defense systems utilizing OLD architecture where an ABC ATPase domain is associated with a TOPRIM nuclease domain. While the ATPase domain is highly conserved, the relative orientation of the TOPRIM nuclease domains and dimerization domains vary between PARIS (PDB ID: 8UX9), Gabija (PDB ID: 8SM3), and the *Thermus scotoductus* Overcoming Lysogenization Defect (*Ts* OLD, PDB ID: 6P74). B) AriB is the predicted effector of PARIS defense. AriB's TOPRIM is homologous to the TOPRIM domains of *Ts* OLD (RMSD 1.1 Å, 20 C-alpha pairs) and GajA (RMSD 1.02 Å, 26 C-alpha pairs). Interestingly, the TOPRIM domain of AriB is found within the DUF4435, which plays a role in AriB dimerization. C) Structural comparison of AriA to Rad50, a universally conserved ATPase, shows a high degree of similarity. AriA primarily differs from Rad50 at Insertion Sequence (IS) 1 (residues 122-180) and IS2 (residues 242-289). IS1 introduces three alpha helices near the nucleotide binding domains of AriA that we hypothesize are involved in trigger recognition. IS2 expands the coiled coil domain with the introduction of two alpha helices which expand the coiled coil domain and enable the homohexameric assembly of AriA subunits in the PARIS complex. D) Structural comparison of PARIS and related OLD systems highlights the unique assembly state of PARIS.

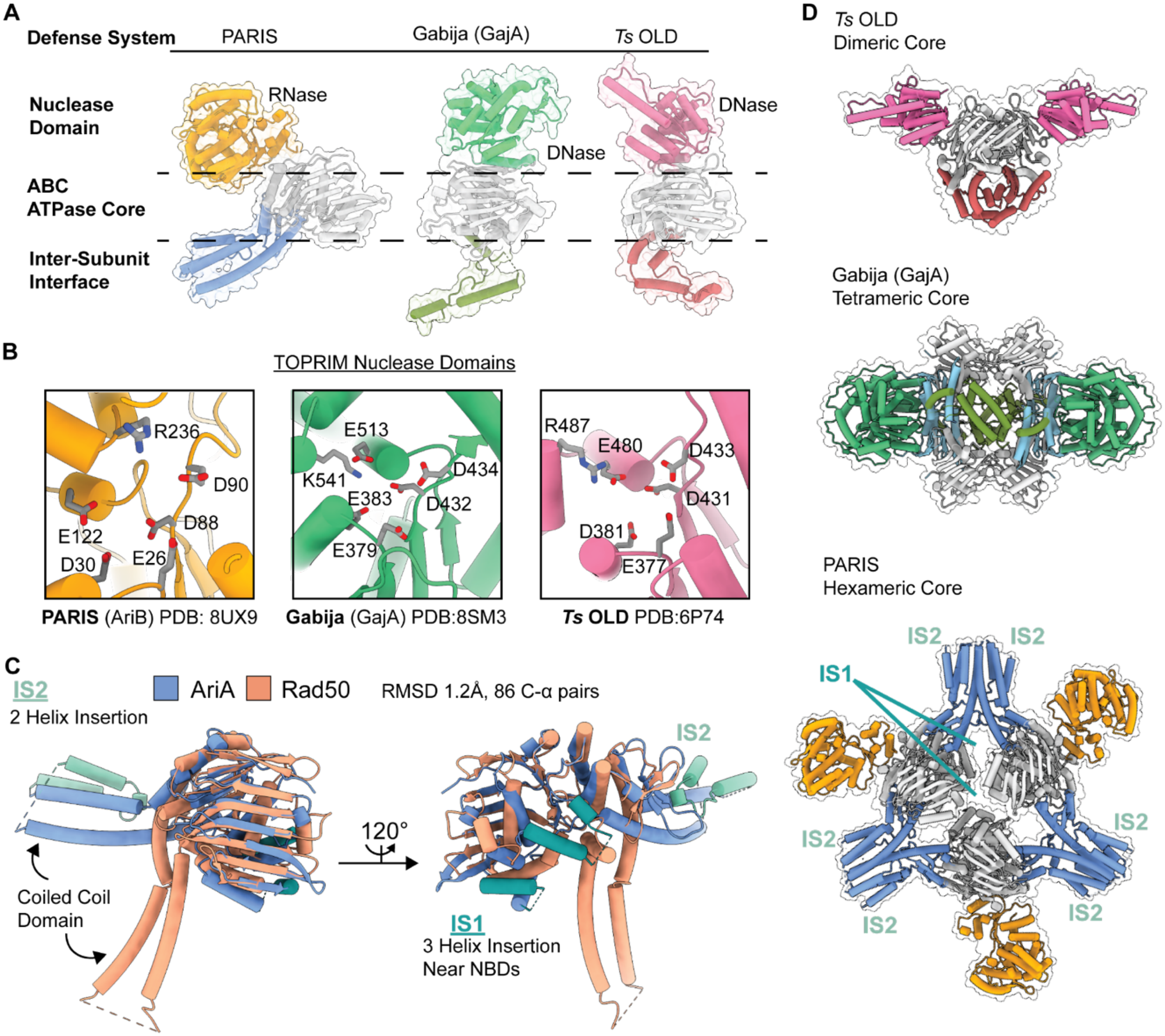

**Supplemental Figure 3: Cryo-EM Workflow for structural determination of the PARIS complex**
Image processing was performed using cryoSPARC v4.3.1. A) After collecting 7,340 movies, micrographs with CTF-fits worse than 8 Å were removed prior to downstream processing (n=5,988). Using a *de novo* template generated from a 200-micrograph subset of this data, 4,078,384 particles were identified and extracted. From 17 selected classes, 1,643,515 particles remained, and a 3-class ab initio reconstruction yielded volumes shown in panel B, particles belonging to junk classes (pink and red) were discarded. C) 25 Representative 2-D Classes from 940,782 particles in panel B. A total of 934,763 particles from 72 classes remained after final 2-D Classification (n=100 classes). D) After successive rounds of *ab initio* and heterogenous refinements, a 532,010 particle volume was obtained, corresponding to the intact PARIS complex with compositional heterogeneity at two AriB attachment sites. Masks were generated for local refinement and 3-D classification to produce the reconstructions shown in E, G, and H, respectively. E) Local Refinement of one asymmetric unit of the PARIS immune complex with 2-D and 3-D FSC data shown below F) Close up of density map and model from E demonstrating map quality G) C3 reconstruction of the PARIS complex at 3.71 Å-resolution with the experimental structure determined in E fit into the density. H) Cryo-EM reconstruction of PARIS with AriB in the 'trans' arrangement at approximately 3.93 Å-resolution with experimental structure determined in E fit into the density.

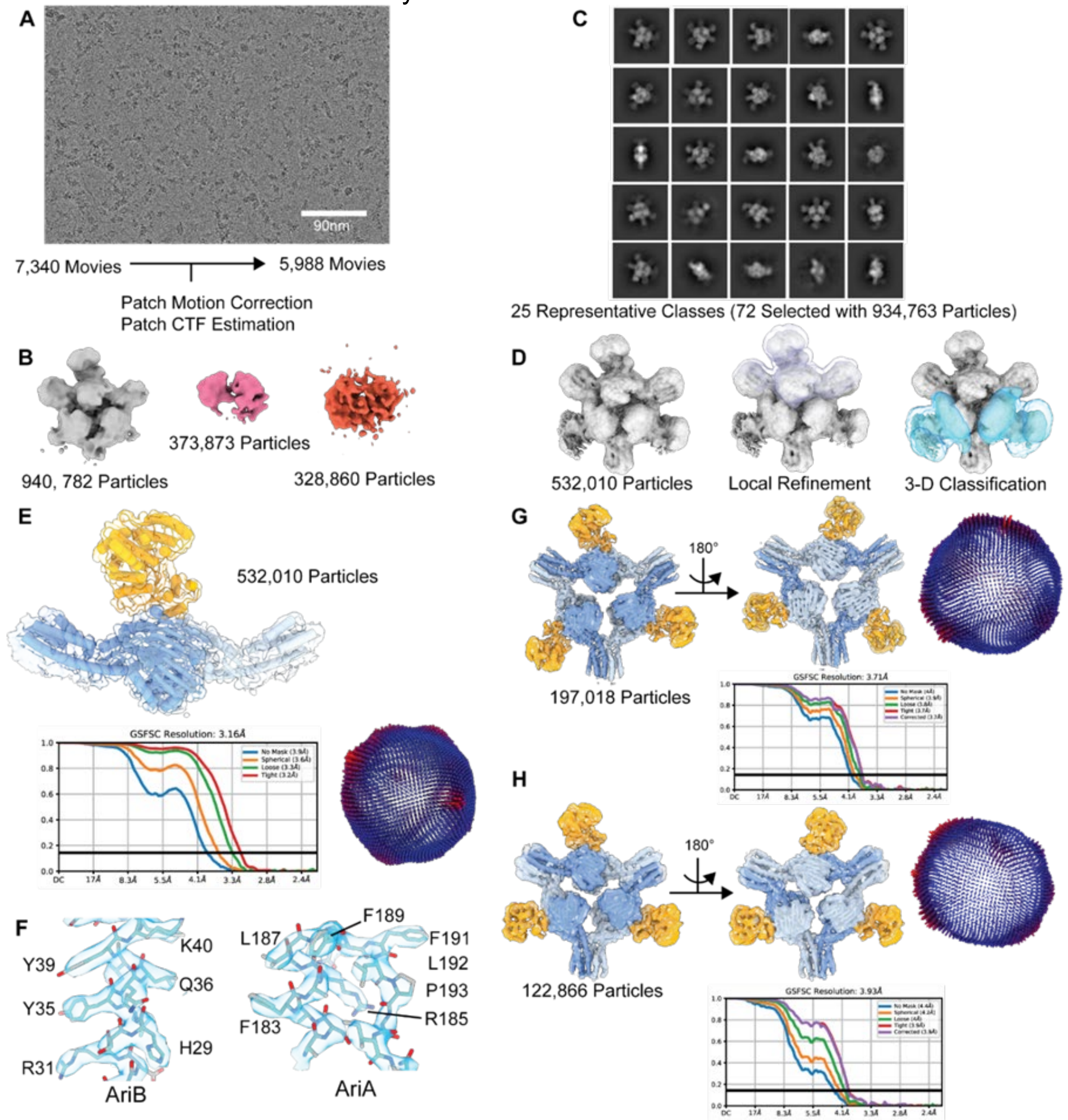

**Supplemental Figure 4: Purification of activated AriB**

A) Size exclusion chromatography calibration curve with estimated masses of AriA-Ocr and activated AriB complexes. B) Introduction of the Strep-tag at the C- or N- terminus of AriA, but on the C-terminus of AriB, interferes with PARIS anti-phage defense. C) Purification of activated AriB after mixing cell lysates containing strep-tagged AriB and non-tagged Ocr.

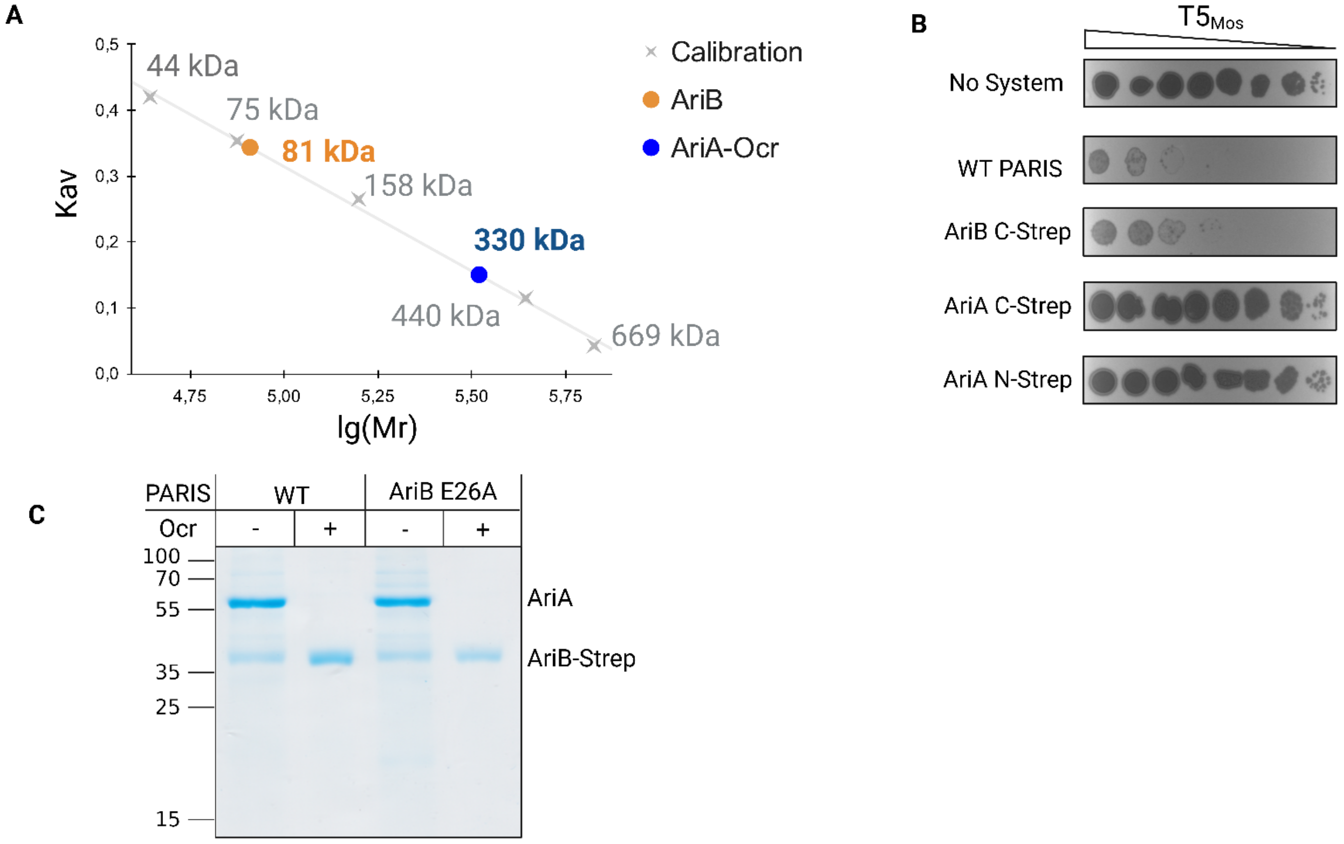

**Supplemental Figure 5: Cryo-EM workflow for Strep-Ocr pulldowns of AriA**

Image processing and analysis was performed in cryoSPARC v4.3.1. A) Representative micrograph from a total of 10,340 movies. Micrographs with CTF-fits worse than 8 Å-resolution were discarded. Particle picking on 9,399 micrographs identified 4,415,782 particles which were extracted and subjected to 2-D Classification. B) Selected 2-D classes corresponding to the AriA hexamer contained 667,479 particles. C) 2-Class *ab initio* reconstruction of particles from panel B. The gray class corresponds to the AriA homohexamer, while particles associated with the orange class were discarded. Density for the ATPase domains is present, but one blade of the propellor is poorly resolved due to the increased flexibility of the AriA scaffold relative to the fully assembled PARIS complex. D) After iterative rounds of sorting the 392,989 particles from panel C, a stack containing 62,732 particle images was isolated and C3 refinement produced a 4.4 Å-resolution reconstruction of the AriA scaffold.

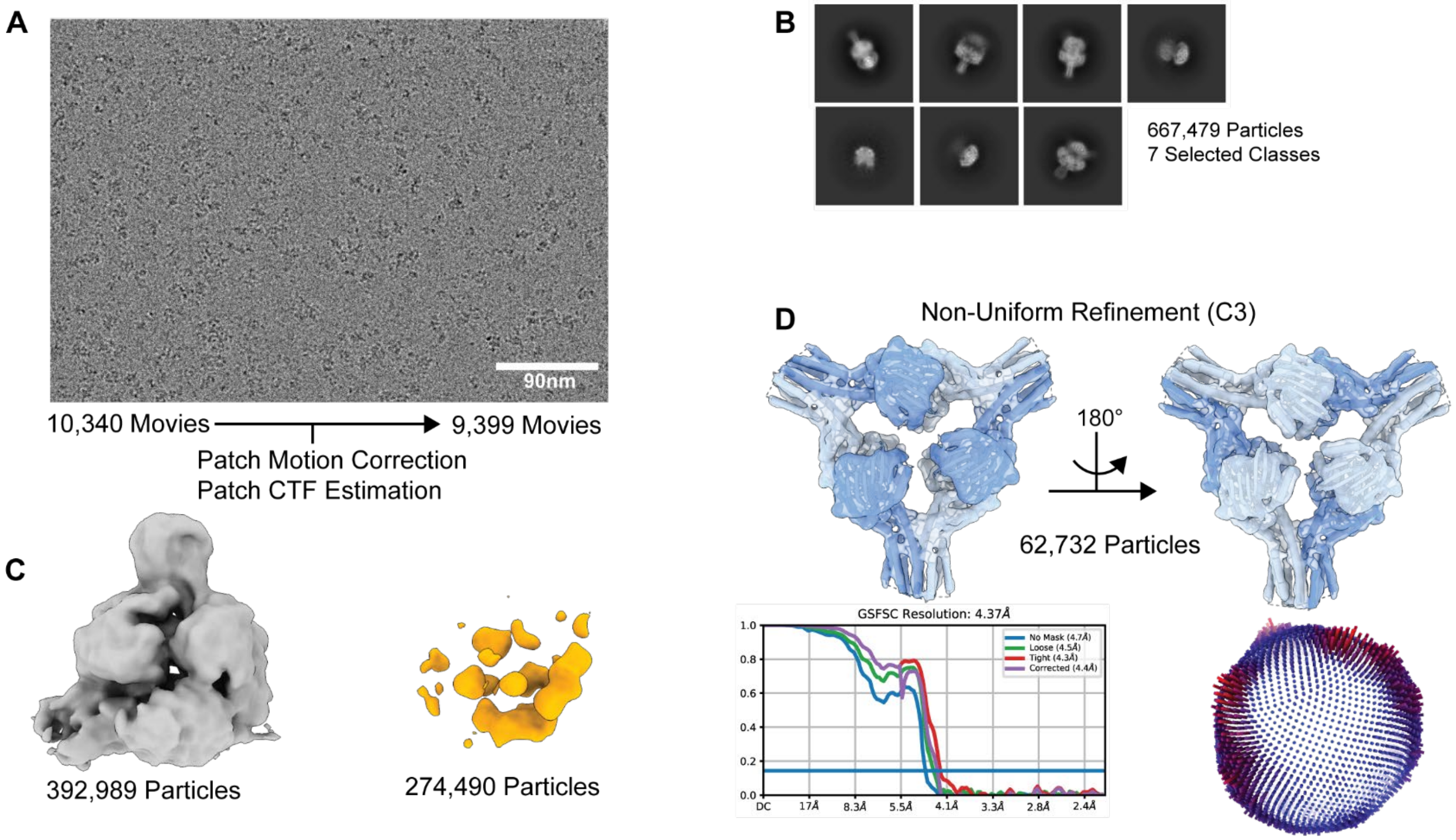

**Supplemental Figure 6: AriA ATPase activity assays**

A) PARIS mediated hydrolysis of $\alpha^{32}P$ ATP with PARIS alone or with 10-fold excess trigger (T7 Ocr). Reactions were run from 1 to 32 minutes and products were resolved via thin layer chromatography. Reactions with $\alpha^{32}P$ ATP incubated with T4 polynucleotide kinase were used as a positive control. Images are representative of reactions that were performed in triplicate. B) Ocr alone was incubated with $\alpha^{32}P$ ATP for 32 minutes indicating that the trigger alone does not hydrolyze ATP. C) Concentration of ADP formed at each time point was quantified and plotted. Data points represent means of three experiments with standard deviation shown.

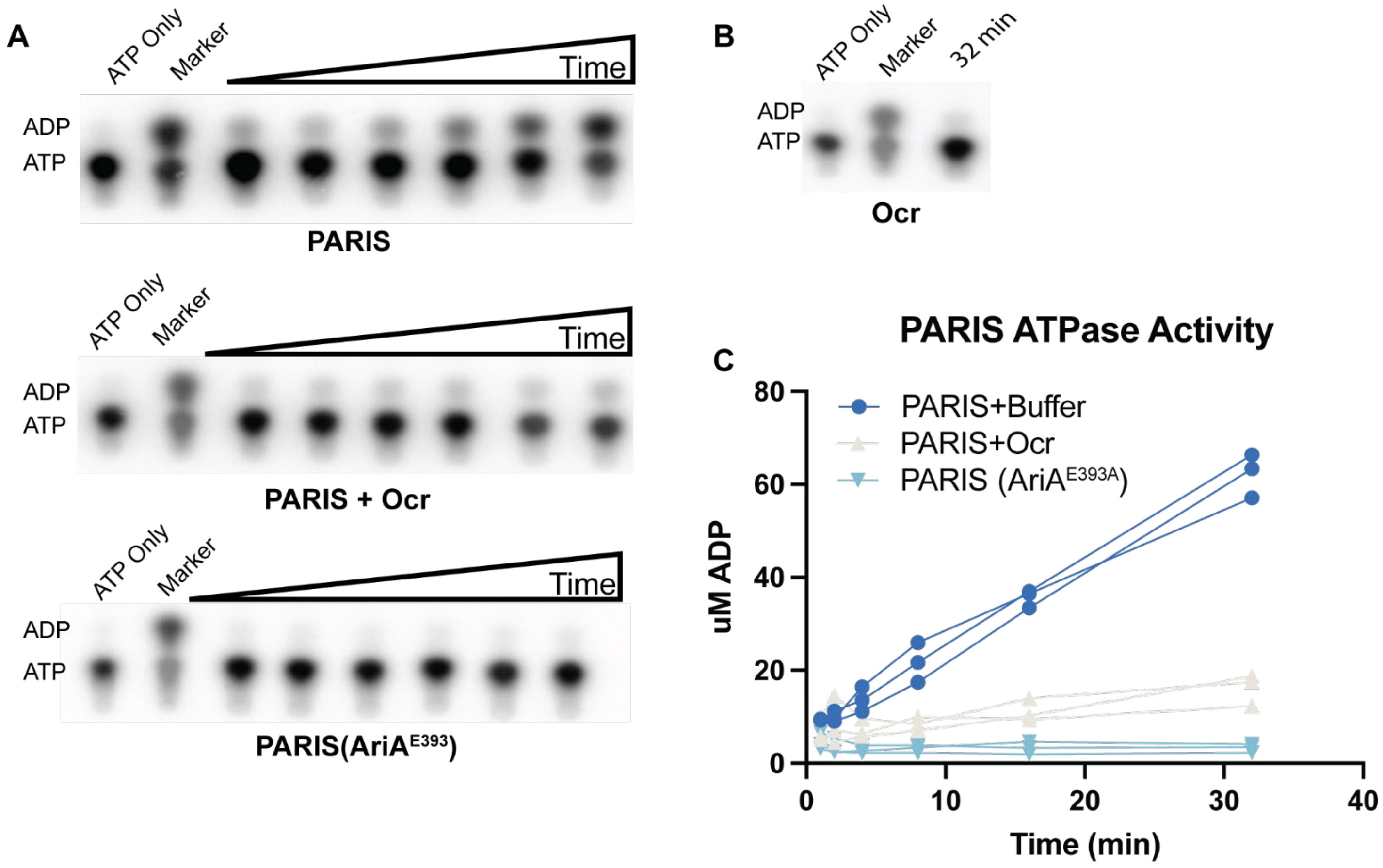

**Supplemental Figure 7: PARIS activation does not result in total RNA or DNA degradation.**

A) Total DNA was extracted from *E. coli* MG1655 carrying plasmids pFR85 (PARIS) and pFD250 ($P_{PhlF}$-Ocr) at different time points after induction Ocr with or without DAPG (n=3), or from a non-induced control. Samples were run on a 1% agarose TAE gel. B) Total RNA was extracted from *E. coli* MG1655 carrying plasmid pFR85 (PARIS) or pFR66 (sfGFP), and pFD250 ($P_{PhlF}$-Ocr) at different time points after induction of Ocr expression with DAPG. Samples were run on a TBE Urea (7M) acrylamide (10%) gel and stained with SYBR-Gold. A representation of 3 replicates is shown. C) TUNEL assay, demonstrating the lack of accumulation of dsDNA breaks in $PARIS^+$ cells 2 hours after Ocr induction, as measured through terminal deoxynucleotidyl transferase labeling of free 3'-OH groups in DNA. As a positive control of DNA damage, cells were treated with 0.1% $H_2O_2$.

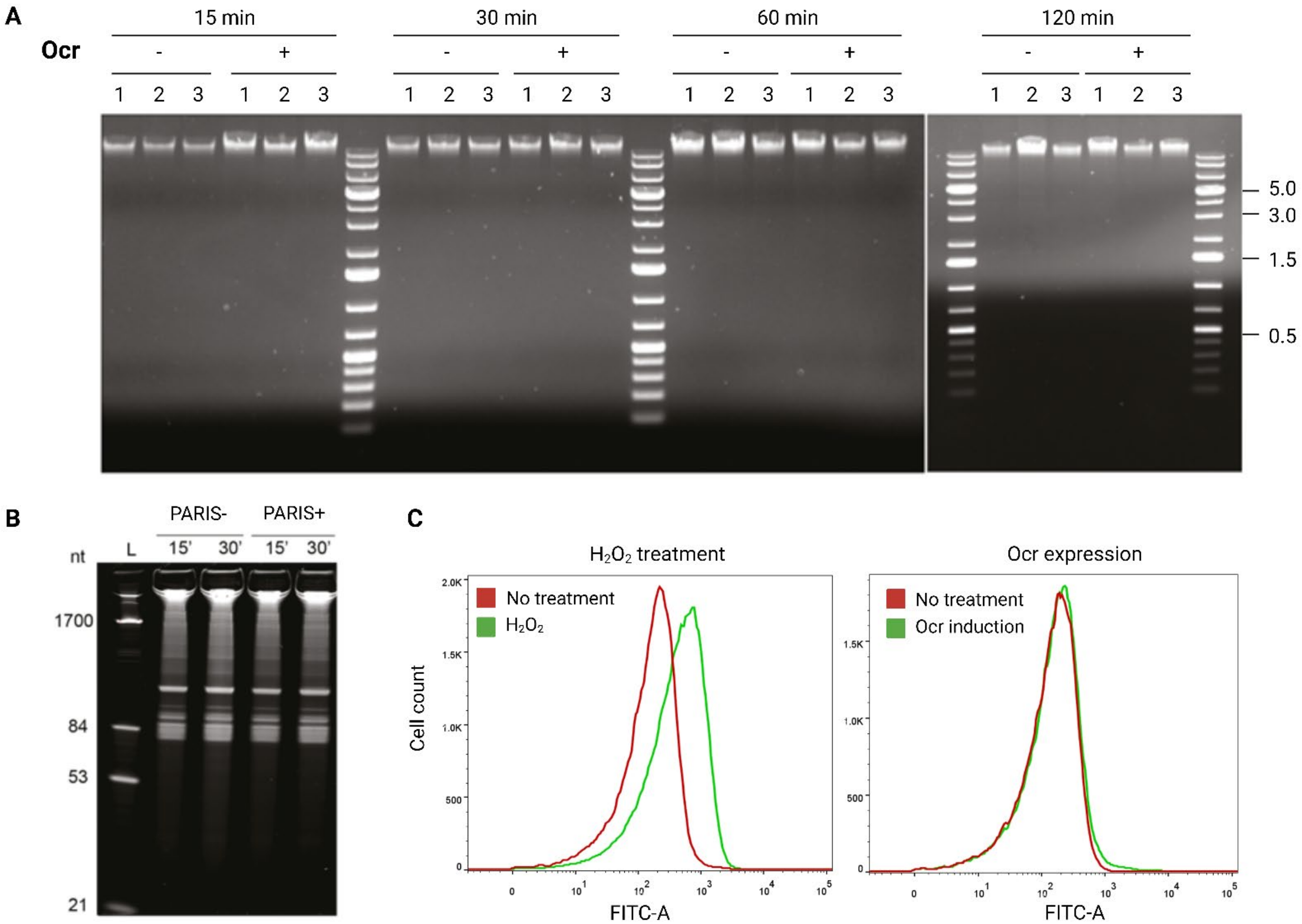

**Supplemental Figure 8: PARIS activation results in DNA compactization akin to inhibition of translation with chloramphenicol.**

A) Hi-C contact maps (bin: 4kb). From left to right: WT (pFR66 + pFD245 control vectors), PARIS (pFR85) + $P_{PhlF}$-ocr (pFD250), and chloramphenicol treated cells. Top and bottom rows correspond to 15 and 30 minutes after induction with DAPG or treatment with chloramphenicol. B) Ratio between contact maps of WT at t = 30 min and contact maps 30 min after either PARIS activation (left) or chloramphenicol treatment (right).

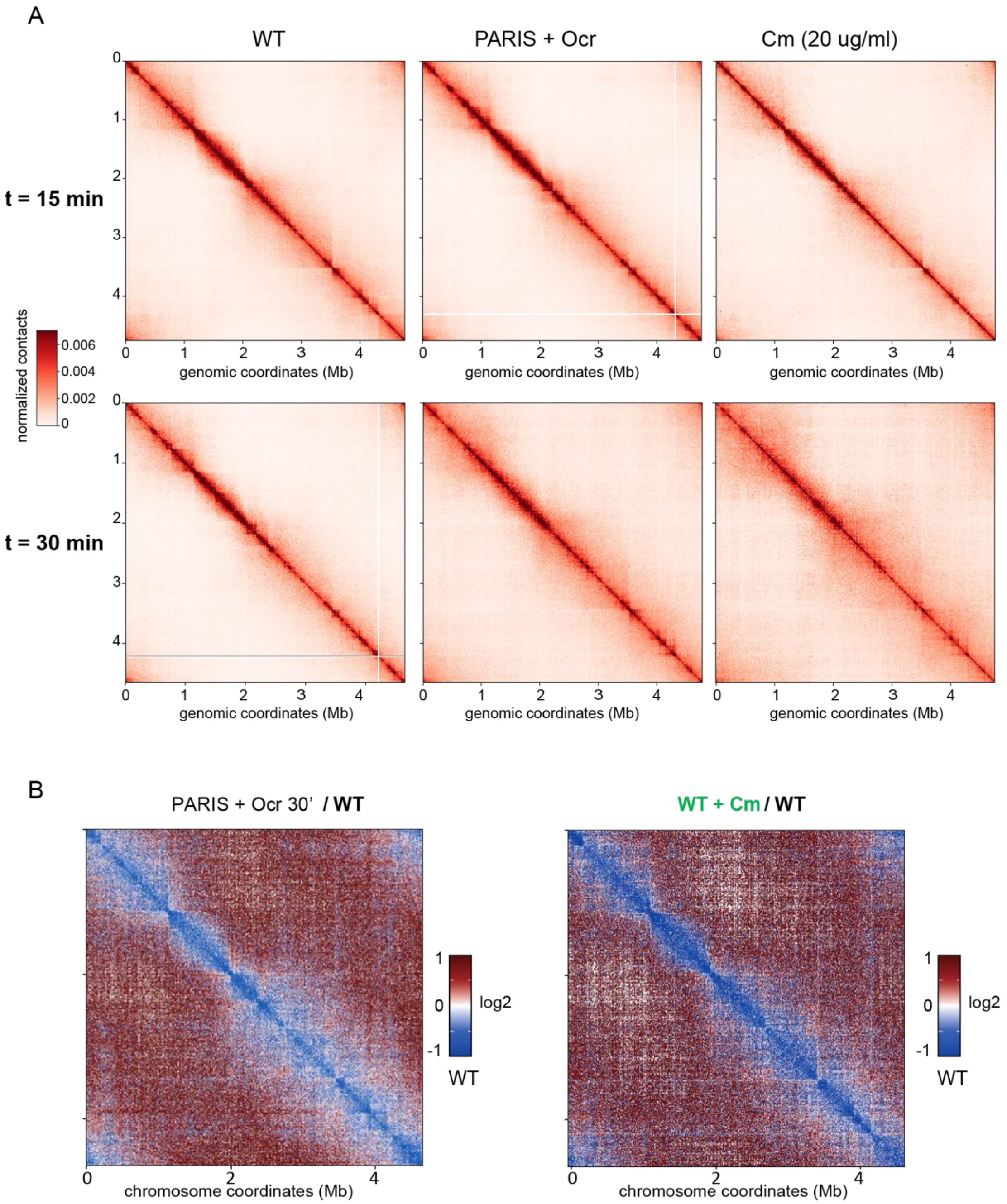

**Supplemental Figure 9: Identification of the T5 genomic region required for productive infection of PARIS$^+$ cells**

A) Growth of PARIS$^-$ or PARIS$^+$ cells infected with PARIS-sensitive phage $T5_{Mos}$ at low or high MOI. Phage was added at t=0. B) Phage T5 variants with non-essential deletions in the region encoding tRNAs are sensitive to PARIS defense. Boundaries of the deletions are shown on the right, tRNA genes are highlighted in red. C) Overexpression of Fragment 1 (31885-32870) derived from the deletion in the phage $T5_{123}$ partially rescues $T5_{123}$ infection of the PARIS$^+$ culture.

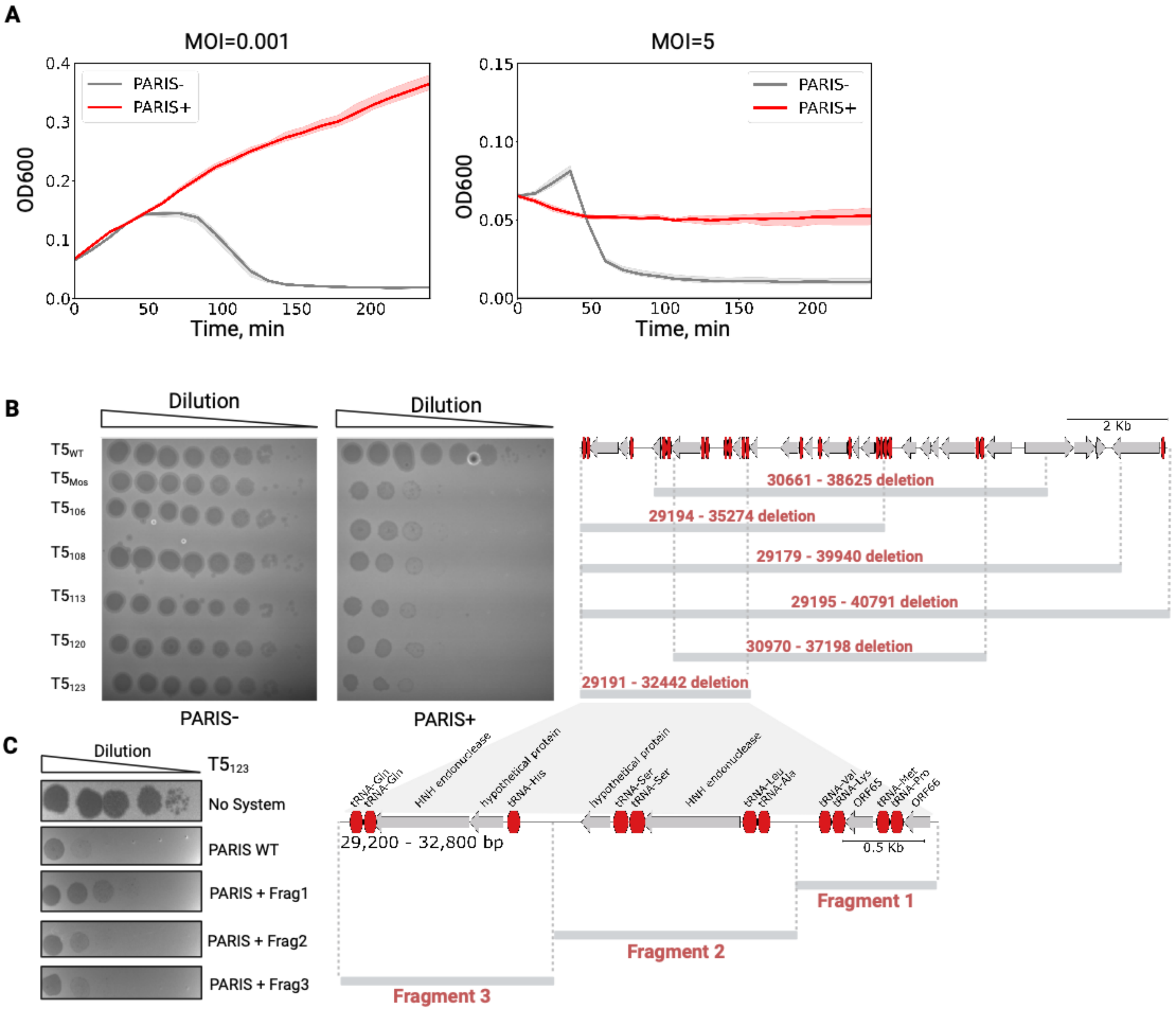

**Supplementary Figure 10. Conservation of AriA and rooted tree of AriB.** A) Alignment of homologs of AriA representative of the different PARIS clades. Different domains of AriA are indicated. Grey scale represents % of identity between homologs. B) Phylogenetic tree of DUF4435 domain of AriB using M5 Ribonuclease (TOPRIM) as an outgroup.

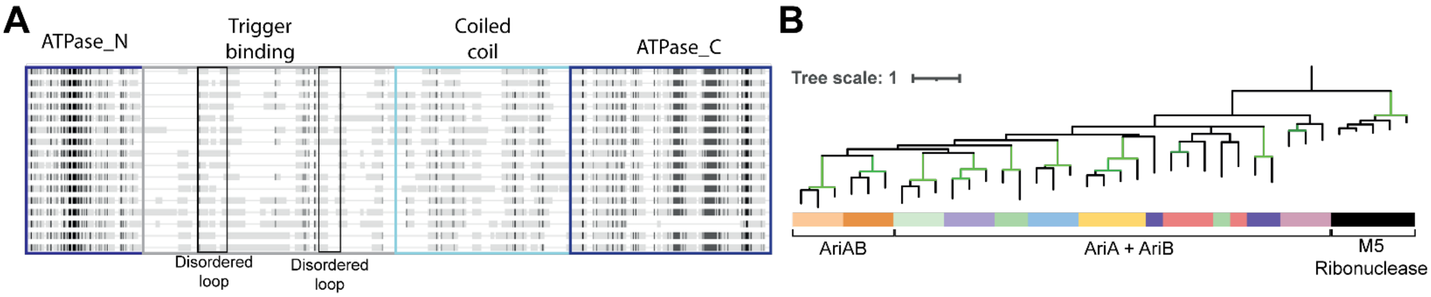
