## Supplementary Table 1 for "Viral proteins activate PARIS-mediated tRNA degradation and viral tRNAs rescue infection"

**Supplementary Table 1. Cryo-EM data collection, refinement, and validation statistics**

| | PARIS Asymmetric Unit (EMDB-42719) (PDB 8UX9) | PARIS 'cis' Arrangement (EMDB-43103) | PARIS 'trans' Arrangement (EMDB-43104) | AriA from Ocr Pulldown (EMDB-43105) |
|---|---|---|---|---|
| **Data collection and processing** | | | | |
| Magnification | 36,000X | 36,000X | 36,000X | 36,000X |
| Voltage (kV) | 200 | 200 | 200 | 200 |
| Electron exposure (e–/Å$^2$) | 56.42 | 56.42 | 56.42 | 56.31 |
| Defocus range (μm) | -0.6, -2.5 | -0.6, -2.5 | -0.6, -2.5 | -0.6, -2.5 |
| Pixel size (Å) | 0.552 | 0.552 | 0.552 | 0.552 |
| Symmetry imposed | C1 | C3 | C1 | C3 |
| Initial particle images (no.) | 4,078,384 | 4,078,384 | 4,078,384 | 667,479 |
| Final particle images (no.) | 532,010 | 197,018 | 122,866 | 62,732 |
| Map resolution (Å) | 3.20 | 3.71 | 3.93 | 4.37 |
| FSC threshold | FSC=0.143 | FSC=0.143 | FSC=0.143 | FSC=0.143 |
| Map resolution range (Å) | 2.64-5.95 | 2.84-6.75 | 2.96-7.53 | 3.72-7.89 |
| **Refinement** | | | | |
| Initial model used (PDB code) | N/A | | | |
| Model resolution (Å) | 3.2 | | | |
| FSC threshold 0.143 | FSC=0.143 | | | |
| Model resolution range (Å) | 2.5-5 | | | |
| Map sharpening *B* factor (Å$^2$) | 117.7 | | | |
| Model composition | | | | |
| Non-hydrogen atoms | 7,783 | | | |
| Protein residues | 1,104 | | | |
| Ligands | AGS:2 | | | |
| *B* factors (Å$^2$) | | | | |
| Protein | 16.3/115.97/56.81 | | | |
| Ligand | 48.29/95.39/72.55 | | | |
| R.m.s. deviations | | | | |
| Bond lengths (Å) | .003(0) | | | |
| Bond angles (°) | .501(0) | | | |
| Validation | | | | |
| MolProbity score | 1.86 | | | |
| Clashscore | 8.56 | | | |
| Poor rotamers (%) | 0.34 | | | |
| Ramachandran plot | | | | |
| Favored (%) | 94.09 | | | |
| Allowed (%) | 5.82 | | | |
| Disallowed (%) | .09 | | | |
