## Supplementary Table 2 for "Viral proteins activate PARIS-mediated tRNA degradation and viral tRNAs rescue infection"

**Supplementary Table S2. Bacterial strains, phages, and plasmids used in the study**

| ***E. coli* strain** | **Comment** | | **Source** |
|---|---|---|---|
| BW25113 | *E. coli* K12 F– Δ(*araD-araB*)567 Δ*lacZ*4787(::*rrnB*-3) λ– *rph*-1 Δ(*rhaD-rhaB*)568 *hsdR*514 | | Lab stock |
| MG1655 | *E. coli* K12 F-, λ-, *ilvG*-, *rfb*-50, *rph*-1 | | Lab stock |
| XL1-Blue | *E. coli* K12 *recA*1 *endA*1 *gyrA*96 *thi*-1 *hsdR*17 *supE*44 *relA*1 *lac* [F'*proAB lacIqZ*ΔM15 Tn10], TetR | | Evrogen |
| **Phage** | **Comment** | | **Source** |
| T7 | | | Lab stock |
| T5 | | | Lab stock |
| T5$_{Mos}$ | T5 with 30661-38625 deletion | | [1] |
| T5$_{106}$ | T5 with 29194-35274 deletion | | |
| T5$_{108}$ | T5 with 29179-39940 deletion | | |
| T5$_{113}$ | T5 with 29195-40791 deletion | | |
| T5$_{120}$ | T5 with 30970-37198 and 24956-26633 deletions | | |
| T5$_{123}$ | T5 with 29191-32442 deletion | | |
| **Plasmid Name** | **Plasmid Map** | **Comment** | **Source** |
| pFR66 | https://benchling.com/s/seq-eyqx3mULGDzw04wc28W7?m=slm-KMqDsXL62sGpcvkRDd08 | sfGFP, Tet promoter, Kan$^R$ | [2] |
| pFR85 (PARIS) | https://benchling.com/s/seq-P0S4QJUYX0D54AVNp81e?m=slm-DWv2SMSVbNQMxjbonDYa | pFR66 with PARIS-2 from *E.coli* B185, native PARIS-2 promoter + Ptet promoter, Kan$^R$ | [2] |
| pFR85 AriB C-strep | https://benchling.com/s/seq-sUB4VGJfrfQd89nt3msu?m=slm-QbAER6Gp1NJsmGBY4D3N | pFR85 with strep-tag II cloned at the C-terminus of AriB, Kan$^R$ | This work |

| | | | |
|---|---|---|---|
| pSol16<br>(pFR85 AriB E26A) | https://benchling.com/s/seq-WftnRXuL4gjKKI7r10Z7?m=slm-I4J4w4IHYg48zyPsvBdU | pFR85 with AriB E26A mutation in TOPRIM domain, $Kan^R$ | [2] |
| pFR85 AriB E26A C-strep | https://benchling.com/s/seq-OoMBvkn7Mr6FILD4XNgr?m=slm-ElvGNMgAUIA3YfwqTO9u | pFR85 AriB E26A with strep-tag II cloned at the C-terminus of AriB, $Kan^R$ | This work |
| pSol12<br>(pFR85 AriA K39A) | https://benchling.com/s/seq-BKNrelnSgCdOeAOFZlax?m=slm-1snqLus07G5RW3pKWFII | pFR85 with AriA K39A mutation in Walker A ATPase motif, $Kan^R$ | [2] |
| pFR85 AriB C-strep AriA K39A | https://benchling.com/s/seq-1EuCP99koHh69XYbckwh?m=slm-6S1Ckf9EtlmjVicwcPKT | pFR85 AriB C-Strep with AriA K39A mutation in Walker A ATPase motif, $Kan^R$ | This work |
| pFR85 AriB E26A AriA R116E/R119E | https://benchling.com/s/seq-o5OL7dZIrsuUnWBbaVX2?m=slm-9JcE2E6FFj28hC3OMViy | pFR85 AriB E26A with AriA R116E/R119E double mutation in central pore, $Kan^R$ | This work |
| pFR85 AriB E26A C-strep AriA R116E/R119E | https://benchling.com/s/seq-VE3RNuSNCTiPYiJqTYNj?m=slm-TOVPtfnvGWfHGfypKY3A | pFR85 AriB E26A C-strep with AriA R116E/R119E double mutation in central pore, $Kan^R$ | This work |
| pRAW-464 | https://benchling.com/s/seq-WJDrKW8L3zayPXdI2ODo?m=slm-mOeMAJe57kDbC2CEN9BV | pRSFDuet-1 with PARIS-2 (*ariA* & *ariB* from *E. coli* B185 under T7 promoter) with strep-tag cloned at C-terminus of AriB, $Kn^R$ | This work |
| pAG-14 | https://benchling.com/s/seq-OGitMlIvIoTlpTeVW1qh?m=slm-p7zmhMZRoQwJbLFcFsVM | pRSFDuet-1 with PARIS-2 (*ariA* & *ariB* from *E. coli* B185 under T7 promoter) AriB E26A mutation in TOPRIM domain, $Kn^R$ | This work |
| | | pRSFDuet-1 with PARIS-2 (*ariA* & *ariB* from *E. coli* B185 under T7 promoter) AriA E393A mutation in ATPase domain, $Kn^R$ | This work |

| | | | |
|---|---|---|---|
| pT7-Fluc | https://benchling.com/s/seq-2wtCLqR3DGsVXZWhvU4G?m=slm-WEbdjqe6kiooqPfwdM1D | Firefly luciferase gene (*fluc*) cloned under control of T7 promoter, $Amp^R$ | A gift from Dr. Sergiev |
| pBAD Ocr | https://benchling.com/s/seq-Mcb9hM3HgRdwZo0A3YP2?m=slm-2UNmUVUYQHtIqvg3LuOL | Phage T7 Ocr cloned under control of *araBAD* promoter, $Amp^R$ | [6] |
| pBAD Ocr C-Strep | https://benchling.com/s/seq-qECZj1604QHIwJGIK9XJ?m=slm-e7OSyufN44pgTS6RvpIT | pBAD Ocr with strep-tag II cloned at the C-terminus of Ocr, $Amp^R$ | [6] |
| pBAD T5-ORF094 | https://benchling.com/s/seq-luQGd3suHCjMDMMNlRz4?m=slm-SlNU3D9NRKmvQAe3dmGp | Phage T5 Ptr1 (ORF094) cloned under control of *araBAD* promoter, $Amp^R$ | This work |
| pBAD T5-ORF103 | https://benchling.com/s/seq-MCvxxnzaSNvA9vKhqUsF?m=slm-F6rI9hH0RgcwinY1Hhqs | Phage T5 Ptr2 (ORF103) cloned under control of *araBAD* promoter, $Amp^R$ | This work |
| pBAD T5_123_frag1 | https://benchling.com/s/seq-TDfBPYerlxgpUzxTgUWM?m=slm-tTXMo5qzOHRS2PyIfZGa | Fragment #1 (31885-32870) of the deletion found in $T5_{123}$ cloned under control of *araBAD* promoter, $Amp^R$ | This work |
| pBAD T5_123_frag2 | https://benchling.com/s/seq-0f9arhOZSmPh2UIVFkAw?m=slm-7azJAuH418UYDTXaaD46 | Fragment #2 (30456-31884) of the deletion found in $T5_{123}$ cloned under control of *araBAD* promoter, $Amp^R$ | This work |
| pBAD T5_123_frag3 | https://benchling.com/s/seq-zbxfbPNLcJyAiRfuKFT2?m=slm-YHWQhlE8xpywqjU5XFv7M | Fragment #3 (29203-30455) of the deletion found in $T5_{123}$ cloned under control of *araBAD* promoter, $Amp^R$ | This work |
| pBAD *E.coli* $tRNA^{Lys}$ | https://benchling.com/s/seq-PBfQldqJYlgCeL11vHOq?m=slm-GesuVCZ1c4UPdrwixsVG | *E.coli* $tRNA^{Lys}$ (UUU) cloned together with 20 bp upstream and downstream under control of *araBAD* promoter, $Amp^R$ | This work |

| | | | |
|---|---|---|---|
| pBAD T5 tRNA$^{Lys}$ | https://benchling.com/s/seq-vMWoZV7cTRTJmxHolzoL?m=slm-lwz3eDeiC18WokES0QGM | T5 tRNA$^{Lys}$ (UUU) cloned together with 20 bp upstream and downstream under control of *araBAD* promoter, Amp$^{R}$ | This work |
| pBAD T5 ORF65 | https://benchling.com/s/seq-kepEtQyKZHSpfPTBXnZY?m=slm-aE9t7dx2o22kvxnr0br4 | Phage T5 ORF065 cloned under control of *araBAD* promoter, Amp$^{R}$ | This work |
| pBAD T5 ORF66 | https://benchling.com/s/seq-8UTEgAmBE0Ak0Nn2SyHs?m=slm-h2ZccJmYur0Uzi51Uk9U | Phage T5 ORF066 cloned under control of *araBAD* promoter, Amp$^{R}$ | This work |
| pBAD T5 tRNA$^{fMet}$ | https://benchling.com/s/seq-LfKR2YCCL2qQiZf22DBo?m=slm-CLh9Vitt22QkOpqcIJbm | T5 tRNA$^{fMet}$ (CAU) cloned together with 20 bp upstream and downstream under control of *araBAD* promoter, Amp$^{R}$ | This work |
| pBAD T5 tRNA$^{Pro}$ | https://benchling.com/s/seq-Myxb3aGPLIU0aY5A6s17?m=slm-vJzCOuUPlrka2cLyxp1k | T5 tRNA$^{Pro}$ (UGG) cloned together with 20 bp upstream and downstream under control of *araBAD* promoter, Amp$^{R}$ | This work |
| pBAD T5 tRNA$^{Val}$ | https://benchling.com/s/seq-RtF7XESubeGYeh21XQ1i?m=slm-H1QDnkm5B94iE16Nopvn | T5 tRNA$^{Val}$ (UAC) cloned together with 20 bp upstream and downstream under control of *araBAD* promoter, Amp$^{R}$ | This work |
| pBAD *E.coli* tRNA$^{Lys}$ mut1 | https://benchling.com/s/seq-lYEcOBedbkp0FTsFySax?m=slm-XPkoZgQVNSNHZU93axVR | pBAD *E.coli* tRNA$^{Lys}$ with mutations in the anticodon stem-loop mimicking T5 tRNA$^{Lys}$ (31A->U, 39U->A), Amp$^{R}$ | This work |
| pBAD *E.coli* tRNA$^{Lys}$ mut2 | https://benchling.com/s/seq-dvmea4IbqeqorFmoBUOA?m=slm-JOeMISpTgEnySwizxegx | pBAD *E.coli* tRNA$^{Lys}$ with mutations in the anticodon stem-loop mimicking T5 tRNA$^{Lys}$ (29U->G, 41A->C), Amp$^{R}$ | This work |
| pAG-8 | ???? | pRSFDuet-1 with His6-TwinStrep-SUMO at the N-terminus of Ocr, T7 promoter, Kn$^{R}$ | This work |

| | | | |
|---|---|---|---|
| pAG-11 | https://benchling.com/s/seq-n2KnqpVRlv8x4lo4NEWk?m=slm-rNrKpvYLgs1HJlslvuIV | pET-Duet with Ocr carrying Strep-tag on C-terminus, T7 promoter, $Amp^R$ | This work |
| pRAW-401 | https://benchling.com/s/seq-CoC6yFNJ755I6Pp32O2l?m=slm-NOM1jk1TNNivclNE8nXd | Vector backbone that has His6-TwinStrep-SUMO | [4] |
| pFD200 | https://benchling.com/s/seq-rUkLXHvua674pynDfnm5?m=slm-8YQXlfp0QQ1Xu8necS3s | pFR56 with dCas9 *ccdB* cloned under the control of $P_{PhlF}$ promoter, $Cm^R$ | This work |
| pFR94 | https://benchling.com/s/seq-pitmlcFiBDtqOCLy6jUe?m=slm-ARwDRtOCZuH4gdkL4YtB | pBAD18 with *0.3* gene encoding Ocr under the control of *araBAD* promoter | [2] |
| pFD245 | https://benchling.com/s/seq-IO4LSvihp3CzlnznYfk4?m=slm-EkG77x6ylh1VTUABLQky | pFD200 with *sfGFP* cloned under the control of $P_{PhlF}$ promoter, $Cm^R$ | This work |
| pFD250 | https://benchling.com/s/seq-3Y0xK3fj0qRzXKjSqkrH?m=slm-WXcGNkiVOWzJvLAupoOb | pFD200 with *0.3* gene encoding Ocr under the control of $P_{PhlF}$ promoter, $Cm^R$ | This work |
| pFD287 | https://benchling.com/s/seq-PHnTolybLsHXKBFWv4UH?m=slm-Sny163elYSt2RsiAMsdr | T5 $tRNA^{Lys}$ cloned with 20 bp upstream and under the control of *araBAD* promoter, $Amp^R$ | This work |
| pDB288 | | pFR85 with AriA R116E mutation in central pore, $Kan^R$ | This work |
| pDB289 | | pFR85 with AriA R119E mutation in central pore, $Kan^R$ | This work |
| pDB290 | | pFR85 with AriA R116A mutation in central pore, $Kan^R$ | This work |
| pDB291 | | pFR85 with AriA R119A mutation in central pore, $Kan^R$ | This work |
| pDB292 | | pFR85 with AriA E393A mutation in ATPase domain, $Kan^R$ | This work |

| | | | |
|---|---|---|---|
| pDB293 | | pFR85 with AriB R31E mutation in AriA-interacting interface, $Kan^R$ | This work |
| pDB294 | | pFR85 with AriB E285R mutation in AriB dimerization interface, $Kan^R$ | This work |
| pDB295 | | pFR85 with AriB F137A mutation in AriB dimerization interface, $Kan^R$ | This work |
| pFD290 | | pFR85 with AriA R168A mutation in radial pore, $Kan^R$ | This work |
| pFD292 | | pFR85 with AriA R172A mutation in radial pore, $Kan^R$ | This work |
| pFD293 | | pFR85 with AriA R168E mutation in radial pore, $Kan^R$ | This work |
| pFD295 | | pFR85 with AriA R172E mutation in radial pore, $Kan^R$ | This work |
| pFD296 | | pFR85 with AriB R28E mutation in AriA-interacting interface, $Kan^R$ | This work |
| pFD297 | | pFR85 with AriB R28E R31E mutations in AriA-interacting interface, $Kan^R$ | This work |

**Supplementary Table S2. Primers used for plasmids cloning**

| **Name** | **Sequence (5′->3′)** | **Template** | **Comment** |
|---|---|---|---|
| pFR66_check_F | GAGCCTTTCGTTTTATTTGATG | pFR85 | Sequencing primers |
| pFR66_check_R | GAGTGATATCCGGAGGCAT | | |
| B185_AriB_Strep_F | CGGGTGGCTCCACTCAATATTTTTCAAAAGCGAG | pFR85 | Introduction of AriB C-strep-tag II to the pFR85 through KLD cloning |
| B185_AriB_Strep_R | CAGTTCGAAAAATGATAATTGGCCAGCCTAG | | |
| B185_AriA_check_R | TAGTGGGCGGCAAATTATC | pFR85 | Sequencing primer |
| AriA_ATPK39A_F | ACCTGCACCATTCTTTC | pFR85 | Introduction of mutations through KLD cloning |
| AriA_ATPK39A_R | GCGTCTACATTAATACATGTTATAGC | | |

| | | | |
|---|---|---|---|
| AriA_R116A_R119 A_F | AGTGAGCTTCACTTTCTATCTCTCTC | pFR85 | |
| AriA_R116A_R119 A_R | TGGCGGAACGAGATGTTAAATC | | |
| AriA_R116E_R119E _F | AGTGTTCTTCACTTTCTATCTCTCTC | pFR85 | |
| AriA_R116E_R119E _R | TGGAAGAACGAGATGTTAAATC | | |
| T5_094_F | TTTGGGCTAACAGGA GGAAGAATTCATGACTTT AAAACCGTATATAGTG | T5 phage DNA | Cloning of T5 PARIS triggers through Gibson Assembly |
| T5_094_R | CAGCCAAGCTTGCGGCCGCGAGCTCTTA GTCATTATTAAACTCCACTG | | |
| T5_103_F | TTTGGGCTAACAGGAGGAAGAATTCGTG AGTAGACTAACTGATTTACTTA | T5 phage DNA | |
| T5_103_R | CAGCCAAGCTTGCGGCCGCGAGCTCTTA CTCCA CCAGGCCA | | |
| pBAD_For | ATGCCATAGCATTTTTATCC | pBAD18 | pBAD sequencing primers |
| pBAD_Rev | GATTTAATCTGTATCAGGCTG | | |
| pBAD256b_F | TTTTTATAACCTCCTTAGAGCTCG | pBAD18 | Amplification of pBAD backbone for Gibson Assembly |
| pBAD256b_R | GTCGACCTGCAGGCATGC | | |
| pBAD256_T5del_12 3 F | AGCTCTAAGGAGGTTATAAAAACGTGGG GTTGGTAGCCTC | T5 phage DNA | Cloning of fragments found in $T5_{123}$ deletion through Gibson Assembly |
| pBAD256_T5del_12 3_1 R | TTGCATGCCTGCAGGTCGACATTCAGTTG CCGCTGAATCTAGTG | | |
| pBAD256_T5del_12 3 R | TTGCATGCCTGCAGGTCGACTACTCAGCG AGGCTTGGAAGG | T5 phage DNA | |
| pBAD256_T5del_12 3_3 F | CTCTAAGGAGGTTATAAAAACCCGACAT GCGTTCGGGGGTA | | |
| pBAD256_T5del_12 3_2 F | CTCTAAGGAGGTTATAAAAAAGTGTATT GGGTTTGCGAGA | | |

| | | | |
|---|---|---|---|
| pBAD256_T5del_123_2 R | TTGCATGCCTGCAGGTCGACAAAGTCTTTCCAAGCGCATC | T5 phage DNA | |
| pBAD256_T5_66_F | AGCTCTAAGGAGGTTATAAAAAGTGAAGTATAAAATTGCATC | T5 phage DNA | Cloning of ORFs found in $T5_{123}$ deletion through Gibson Assembly |
| pBAD256_T5_66_R | TTGCATGCCTGCAGGTCGACTTATTTCTTCAGCCACAAC | | |
| pBAD256_T5_65_F | AGCTCTAAGGAGGTTATAAAAAATGAAAGTATTTCGATTAG | T5 phage DNA | |
| pBAD256_T5_65_R | TTGCATGCCTGCAGGTCGACTTAGTAACCATTATATACTTC | | |
| pBAD256_T5_Pro_F | TTTTTGGGCTAGCGAATTCGAGCTCGTGGCTGAAGAAATAATTTTC | T5 phage DNA | Cloning of tRNAs found in $T5_{123}$ deletion through Gibson Assembly |
| pBAD256_T5_Pro_R | CAAGCTTGCATGCCTGCAGGTCGACAGATCTAACCCGCAATTG | | |
| pBAD256_T5_fMet_F | TTTTTGGGCTAGCGAATTCGAGCTCTTCATTGGAGACCAAAC | T5 phage DNA | |
| pBAD256_T5_fMet_R | CAAGCTTGCATGCCTGCAGGTCGACTTCGAAATCCTCTATTAAAATG | | |
| pBAD256_T5_Lys_F | TTTTTGGGCTAGCGAATTCGAGCTCAAGTATATAATGGTTACTAAGGG | T5 phage DNA | |
| pBAD256_T5_Lys_R | CAAGCTTGCATGCCTGCAGGTCGACATACTAACCGAGCAAAAATTTG | | |
| pBAD256_T5_Val_F | TTTTTGGGCTAGCGAATTCGAGCTCGTGCAACCCACCAAATTTTTG | T5 phage DNA | |
| pBAD256_T5_Val_R | CAAGCTTGCATGCCTGCAGGTCGACTTTTTGAAATTTCTTGGACTTGG | | |
| pBAD256_T5_LysColi_F | TTTTTGGGCTAGCGAATTCGAGCTCACCGAAAGCAACGAAAAAGTG | BW25113 genomic DNA | Cloning of *E. coli* $tRNA^{Lys}$ through Gibson Assembly |
| pBAD256_T5_LystColi_R | CAAGCTTGCATGCCTGCAGGTCGACACCGCTTCCACCTTAGCG | | |
| pBAD_tRNALysE.coli_mut1_F | TAAACAATTGGTCGCAGGTT | | Mutagenesis of *E. coli* $tRNA^{Lys}$ |

| | | | |
|---|---|---|---|
| pBAD_tRNALysE.coli_mut1_R | AAAGACAACTGCTCTACCAACT | pBAD *E.coli* $tRNA^{Lys}$ | through KLD reaction |
| pBAD_tRNALysE.coli_mut2_F | TAATCCATTGGTCGCAGGTT | pBAD *E.coli* $tRNA^{Lys}$ | |
| pBAD_tRNALysE.coli_mut2_R | AAAGTCCACTGCTCTACCAACTG | | |
| F507-F | CAAATAAAACGAAAGGCGTCGGGTAGCACCAGAAGTCTATAG | pFD200 | Cloning of sfGFP cassette using Gibson assembly leading to **pFD245** |
| F508-R | GCCTCCGGATATCACTTTTGCCACCTGCACTCGTCCTTTG | | |
| F509-F | GTGCAGGTGGCAAAAGTGATATCCGGAGGCATATCAAATGAC | pFR66 | |
| F510-R | CTGGTGCTACCCGACGCCTTTCGTTTTATTTGATGCCTGG | | |
| F511-F | GTCGGGTAGCACCAGAAGTCTATAG | pFD200 | Cloning of ocr trigger using Gibson assembly leading to **pFD250** |
| F512-R | TTTTGCCACCTGCACTCGTCC | | |
| F523-F | CATATCAAAGGACGAGTGCAGGTGGCAAAAATGGCTATGTCTAACATGACTTAC | pFR94 | |
| F524-R | CATGATACGAAACGTACCGTATCGTTAAGGTCGAGATTTACTCAGGAGGTTAATTTTAATGGATATTAATACTGAAACTGAGATC | | |
| V318-F | CGGGTATGGAGAAACAGTAGAGAG | pBAD18 | Cloning of T5 $tRNA^{Lys}$ using Gibson assembly leading to **pFD287** |
| F593-R | CAAGCTTGGCTGTTTTGGC | | |
| F617-F | TCGCAACTCTCTACTGTTTCTCCATACCCGAAGTATATAATGGTTACTAAGGGTTGCTAG | T5 phage DNA | |
| F605-R | CTTCTCTCATCCGCCAAAACAGCCAAGCTTGAAATTTGGTGGGTTGCACAGGACTC | | |
| Sol35-F | ATGGTGCAGGTGCATCTACATTAATACATG | pFR85 | Mutation in AriA-K39A of pFR85 (PARIS) using Gibson assembly leading to **pSol12** |
| LC191-R | GTCTAGGGCGGCGGATTTG | | |
| LC192-F | CGCTCTCCTGAGTAGGACAAAT | pFR85 | |
| Sol36-R | TATTAATGTAGATGCACCTGCACCATTCTT | | |

| | | | |
|---|---|---|---|
| Sol41-F | CGTCTGTTAGTTGCGGGGAGGCATGACCGT | pFR85 | Mutation in *ariB*-E26A of pFR85 (PARIS) using Gibson assembly leading to **pSol16** |
| LC191-R | GTCTAGGGCGGCGGATTTG | | |
| LC192-F | CGCTCTCCTGAGTAGGACAAAT | pFR85 | |
| Sol42-R | GTCATGCCTCCCCGCAACTAACAGACGCTT | | |
| B765-F | GAGATAGAAAGTGAAGAACACTTGAGGGAACGAGATGTTA | pFR85 | Mutation in *ariA*-R116E of pFR85 (PARIS) using Gibson assembly leading to **pDB288** |
| LC327-R | CGCCTTCTTGACGAGTTCTT | | |
| TG99-F | GGATCTATCAACAGGAGTC | pFR85 | |
| B766-R | CTCGTTCCCTCAAGTGTTCTTCACTTTCTATCTCTCTCAC | | |
| B767-F | GAACGTCACTTGGAAGAACGAGATGTTAAATCCATGCTTG | pFR85 | Mutation in *ariA*-R119E of pFR85 (PARIS) using Gibson assembly leading to **pDB289** |
| LC327-R | CGCCTTCTTGACGAGTTCTT | | |
| TG99-F | GGATCTATCAACAGGAGTC | pFR85 | |
| B768-R | ATGGATTTAACATCTCGTTCTTCCAAGTGACGTTCACTTTC | | |
| B769-F | GAGATAGAAAGTGAAGCGCACTTGAGGGAACGAGATGTTA | pFR85 | Mutation in *ariA*-R116A of pFR85 (PARIS) using Gibson assembly leading to **pDB290** |
| LC327-R | CGCCTTCTTGACGAGTTCTT | | |
| TG99-F | GGATCTATCAACAGGAGTC | pFR85 | |
| B770-R | CTCGTTCCCTCAAGTGCGCTTCACTTTCTATCTCTCTCAC | | |

| | | | |
|---|---|---|---|
| B771-F | GAACGTCACTTGGCGGAACGAGATGTTA AATCCATGCTTG | pFR85 | Mutation in *ariA*-R119A of pFR85 (PARIS) using Gibson assembly leading to **pDB291** |
| LC327-R | CGCCTTCTTGACGAGTTCTT | | |
| TG99-F | GGATCTATCAACAGGAGTC | pFR85 | |
| B772-R | ATGGATTTAACATCTCGTTCCGCCAAGTG ACGTTCACTTTC | | |
| B773-F | TGCTAATTGATGCGCCAGAGATTTCATTA CATATCGATTG | pFR85 | Mutation in *ariA*-E393A of pFR85 (PARIS) using Gibson assembly leading to **pDB292** |
| LC327-R | CGCCTTCTTGACGAGTTCTT | | |
| TG99-F | GGATCTATCAACAGGAGTC | pFR85 | |
| B774-R | GATATGTAATGAAATCTCTGGCGCATCA ATTAGCACAATTG | | |
| B777-F | TAACAACCGCTTTAGCTCGCAGTTGGGTT AGGAAAATTGG | pFR85 | Mutation in *ariB*-E285R of pFR85 (PARIS) using Gibson assembly leading to **pDB294** |
| LC327-R | CGCCTTCTTGACGAGTTCTT | | |
| TG99-F | GGATCTATCAACAGGAGTC | pFR85 | |
| B778-R | TTTCCTAACCCAACTGCGAGCTAAAGCG GTTGTTAATGAT | | |
| B779-F | AATAGATGCATTTCAGGCGCTCTCACCTT CGGAATATAAATATAAG | pFR85 | Mutation in *ariB*-F137A of pFR85 (PARIS) using Gibson assembly leading to **pDB295** |
| LC327-R | CGCCTTCTTGACGAGTTCTT | | |
| TG99-F | GGATCTATCAACAGGAGTC | pFR85 | |
| B780-R | TATTCCGAAGGTGAGAGCGCCTGAAATG CATCTATTATTATAG | | |

| | | | |
|---|---|---|---|
| F633-F | GATGTAGGCTATGAGGCGAGAGTTATTC GCTCATCATTTTATAACC | pFR85 | Mutation in *ariA*-R168A of pFR85 (PARIS) using Gibson assembly leading to **pFD290** |
| LC327-R | CGCCTTCTTGACGAGTTCTT | | |
| TG99-F | GGATCTATCAACAGGAGTC | pFR85 | |
| F634-R | GAGCGAATAACTCTCGCCTCATAGCCTAC ATCAGAAGATGAAC | | |
| F637-F | GAGCGAAGAGTTATTGCGTCATCATTTTA TAACCGTAAAGCCAGTG | pFR85 | Mutation in *ariA*-R172A of pFR85 (PARIS) using Gibson assembly leading to **pFD292** |
| LC327-R | CGCCTTCTTGACGAGTTCTT | | |
| TG99-F | GGATCTATCAACAGGAGTC | pFR85 | |
| F638-R | CTTTACGGTTATAAAATGATGACGCAATA ACTCTTCGCTCATAGCCTACATCAG | | |
| F639-F | GATGTAGGCTATGAGGAAAGAGTTATTC GCTCATCATTTTATAAC | pFR85 | Mutation in *ariA*-R168E of pFR85 (PARIS) using Gibson assembly leading to **pFD293** |
| LC327-R | CGCCTTCTTGACGAGTTCTT | | |
| TG99-F | GGATCTATCAACAGGAGTC | pFR85 | |
| F640-R | GAGCGAATAACTCTTTCCTCATAGCCTAC ATCAGAAGATGAAC | | |
| F643-F | GAGCGAAGAGTTATTGAATCATCATTTTA TAACCGTAAAGCCAGTG | pFR85 | Mutation in *ariA*-R172E of pFR85 (PARIS) using Gibson assembly leading to **pFD295** |
| LC327-R | CGCCTTCTTGACGAGTTCTT | | |
| TG99-F | GGATCTATCAACAGGAGTC | pFR85 | |
| F644-R | CTTTACGGTTATAAAATGATGATTCAATA ACTCTTCGCTCATAGCCTACATC | | |

| | | | |
|---|---|---|---|
| F645-F | CTGTTAGTTGAAGGGGAACATGACCGTTCGCATCTGTATC | pFR85 | Mutation in *ariB*-R28E of pFR85 (PARIS) using Gibson assembly leading to **pFD296** |
| LC327-R | CGCCTTCTTGACGAGTTCTT | | |
| TG99-F | GGATCTATCAACAGGAGTC | pFR85 | |
| F646-R | GATGCGAACGGTCATGTTCCCCTTCAACTAACAGACGCTT | | |
| B775-F | TTGAAGGGAGGCATGACGAATCGCATCTGTATCAATTAATTTA | pFR85 | Mutation in *ariB*-R31E of pFR85 (PARIS) using Gibson assembly leading to **pDB293** |
| LC327-R | CGCCTTCTTGACGAGTTCTT | | |
| TG99-F | GGATCTATCAACAGGAGTC | pFR85 | |
| B776-R | ATTGATACAGATGCGATTCGTCATGCCTCCCTTCAACTAAC | | |
| F647-F | CTGTTAGTTGAAGGGGAACATGACGAATCGCATCTGTATCAATTAATTTA | pFR85 | Mutation in *ariB*-R28E/R31E of pFR85 (PARIS) using Gibson assembly leading to **pFD297** |
| LC327-R | CGCCTTCTTGACGAGTTCTT | | |
| TG99-F | GGATCTATCAACAGGAGTC | pFR85 | |
| F648-R | ATTGATACAGATGCGATTCGTCATGTTCCCCTTCAACTAACAGACGCTT | | |
| HiFi F - PARIS-2 AriB-strep | GTATAAGAAGGAGATATACATATGAGTTCATGTGCTTAC | pFR85 | Cloning AriB with C-strep-tag leading to **pRAW-464** |
| HiFi R - PARIS-2 AriB-strep | CAGCGGTTTCTTTACCAGACTCGAGTCATTTTTCGAACTGCGG | | |
| HiFi F - PARIS-2 AriA | ACTTTAATAAGGAGATATACCATGGCGATTAGAACAATAAG | pFR85 | Cloning AriA to **pRAW-464** |
| HiFi R - PARIS-2 AriA | TTAAGCATTATGCGGCCGCAAGCTTTCATAAAGATTCATCCTCTTC | | |

| | | | |
|---|---|---|---|
| | TGACTCGAGTCTGGTAAAG | pRAW-464 | Deletion of C-term strep tag from AriB in pRAW-464, for co-expression experiments with strep-tagged trigger |
| | CTCAATATTTTTCAAAAGCGAG | | |
| | TCTGTTAGTTGCGGGGAGGCATG | pRAW-464 | Introduction of mutation in AriB (E26A) |
| | CGCTTTTTAGAGCTCATAG | | |
| | TAATGCTTAAGTCGAACAGAAAG | pRAW-401 | Amplification of vector backbone that has His6-TwinStrep-SUMO leading to **pAG-8** |
| | TCCACCAATCTGTTCTCTG | | |
| | ACAGAGAACAGATTGGTGGAATGGCTATGTCTAACATGAC | pBAD-L18-Ocr-C-Strep | Cloning of T7 Ocr leading to **pAG-8** |
| | TCTGTTCGACTTAAGCATTACTCTTCATCCTCCTCGTAC | | |
| | TTAACCTAGGCTGCTGCC | pET-Duet1 | Amplification of vector backbone of pET-Duet1 leading to **pAG-11** |
| | GGTATATCTCCTTCTTAAAGTTAAACAAAATTATTTC | | |
| | CTTTAAGAAGGAGATATACCATGGCTATGTCTAACATG | pBAD Ocr C-Strep | Cloning of T7 Ocr gene with strep-tag on C-terminus leading to **pAG-11** |
| | GTGGCAGCAGCCTAGGTTAATTATTTTTCGAACTGCGG | | |

### Supplementary Table S3. Nucleotide sequence of the probes used for Northern blot

| Name | Probe | Dye labeled | Sequence 5'-> 3' |
|---|---|---|---|
| **B803** | **$tRNA^{Lys}$** | IRD700 | /5IRD700/GGTGGGTCGTGCAGGATTCGAACCTGCGACCAATTGATTAAA |
| **B806** | **T5 $tRNA^{Lys}$** | IRD800 | /5IRD800/GTTTAAAAGACCAGTGCTCTAAACCAGTTGAGCTAGCAACCC |

| | | | |
|---|---|---|---|
| **B811** | **5S rRNA** | FAM | /56-FAM/TCGGCATGGGGTCAGGTGGGACCACCGCGCTACTGCCGCCAGGCA |
