## Supplementary Movie 1 Caption for "Viral proteins activate PARIS-mediated tRNA degradation and viral tRNAs rescue infection"

**Supplemental Movie 1: Visualization of the PARIS Immune Complex**

Cryo-EM data processing resulted in an initial consensus volume, from which a high-resolution map of one asymmetric unit (AriA2:AriB1) of the PARIS complex was determined assembly. Density for ATPγS in evident in both nucleotide binding domains (NBDs) of the AriA homodimer. A series of electrostatic interactions link AriB to AriA. AriB is located directly above the NDBs of AriA. A structural comparison between the AriB TOPRIM active site and the TOPRIM form Old (Overcoming Lysogenization), reveals spatial conservation of the catalytic residues. From the particle stack used to determine the structure of the asymmetric unit, focused 3-D classification was used to isolate two isomers of the assembled complex, a 'cis' arrangement where all AriB subunits are oriented in the same direction, and a 'trans' arrangement where one subunit of AriB is rotated 180° relative to the others. Viral triggers of PARIS, such as the T7 Ocr protein, release AriB from the complex which forms a homodimeric nuclease that cleaves host tRNALys and induces abortive infection.
