## Supplementary Movie 2 Caption for "Viral proteins activate PARIS-mediated tRNA degradation and viral tRNAs rescue infection"

**Supplemental Movie 2: Ocr induces membrane permeability in cells expressing the PARIS immune system**

Time course video demonstrating PARIS-induced membrane permeability by propidium iodide staining. The cells turn red within 30 minutes of Ocr induction.
